# Nucleotide binding, evolutionary insights and interaction partners of the pseudokinase Unc-51-like kinase 4

**DOI:** 10.1101/2020.06.18.159293

**Authors:** Franziska Preuss, Deep Chatterjee, Sebastian Mathea, Safal Shrestha, Jonathan St-Germain, Manipa Saha, Natarajan Kannan, Brian Raught, Robert Rottapel, Stefan Knapp

**Author notes:** shared first authors (Franziska Preuss, Deep Chatterjee). Correspondence: Stefan Knapp.

## Abstract

Unc-51-like kinase 4 (ULK4) is a pseudokinase that has been linked to the development of several diseases. Even though sequence motifs required for ATP binding in kinases are lacking, ULK4 still tightly binds ATP and the presence of the cofactor is required for structural stability of ULK4. Here we present a high-resolution structure of a ULK4-ATPγS complex revealing a highly unusual ATP binding mode in which the lack of the canonical VAIK motif lysine is compensated by K39, located N-terminal to αC. Evolutionary analysis suggests that degradation of active site motifs in metazoan ULK4 has co-occurred with an ULK4 specific activation loop, which stabilizes the C-helix. In addition, cellular interaction studies using BioID and biochemical validation data revealed high confidence interactors of the pseudokinase and armadillo repeat domains. Many of the identified ULK4 interaction partners were centrosomal and tubulin associated proteins and several active kinases suggesting new roles for ULK4.

**Highlights:** Structure of the ULK4 ATP complex reveals a unique ATP binding mode.

Disease associated mutations modulate ATP binding and ULK4 stability

Degradation of active site motifs co-occurred in evolution with an ULK4 specific activation loop

BioID suggests a role of ULK4 regulating centrosomal and cytoskeletal functions

## Introduction

The human genome encodes about 520 protein kinases. Structural studies revealed that protein kinases share a common topology, the canonical ‘kinase fold’ and large scale sequence comparison have revealed highly conserved amino acid motifs important for kinase catalytic function including (I) the glycine-rich loop, (II) the VAIK motif containing a lysine to bridge the β3 strand to the αC helix in active kinases coordinating the Mg^2+^/triphosphate moiety of the ATP co-factor, (III) the HRD motif with the catalytically indispensable catalytic base, (IV) a conserved asparagine prior to the β7 strand that orients the ATP phosphates and finally (V) the DFG motif at the N-terminus of the activation loop that is important for binding of the phosphate moieties of ATP and Mg^2+^ (Manning et al., 2002). However, about 10% of kinases lack one or more of these conserved motifs, which often renders them catalytically inactive or with significantly reduced catalytic activity (Boudeau et al., 2006; Kwon et al., 2019). These kinases are therefore referred to as pseudokinases. Instead of catalyzing the phosphoryl transfer reaction, their physiological role mediating cellular signaling is scaffolding and to serve as allosteric regulators of active enzymes (recently reviewed by Jacobsen *et al.* (Jacobsen and Murphy, 2017)).

Pseudokinases are present in all major groups of the human kinome and across diverse species (Kwon et al., 2019). The structural features of pseudokinases are accordingly diverse, sharing many regulatory mechanisms observed in canonical kinases (Ha and Boggon, 2018; Jura et al., 2009; Patel et al., 2017; Scheeff et al., 2009; Shrestha et al., 2020; Zeqiraj et al., 2009b). Interestingly, many human pseudokinases have completely lost the ability to interact with nucleotides, while others still bind to and are stabilized by ATP interaction (Murphy et al., 2014). The subgroup of ATP-binding pseudokinases are thought to fulfil their physiological tasks in the nucleotide-bound form. There are several variations of how pseudokinases exert their signaling roles (Murphy et al., 2017). For instance, the transmembrane growth factor receptor HER3 regulates the MAPK and PI3K pathways through interactions with the surface of its pseudokinase C-lobe which constitutes a docking site for client proteins (Citri et al., 2003). When binding, the clients are forced into a specific conformation, resulting in client activation (Jura et al., 2009).

An intriguing example of a complex and dynamic regulatory mechanism mediated by a pseudokinase is Mixed Lineage Kinase domain-Like (MLKL), a key regulator of the non-apoptotic, kinase dependent programmed cell death called necroptosis (Murphy et al., 2013). MLKL comprises a N-terminal 4-helix bundle (4HB) domain and a C-terminal pseudokinase which lacks catalytic activity due to loss of the DFG and HRD motifs. Induction of necroptosis relies on two active kinases, the receptor interacting protein kinase RIPK1 and RIPK3 which interact with MLKL forming the necrosome. Formation of this complex results in phosphorylation of MLKL, triggering a conformational change in MLKL that unleashes the 4HB domain resulting in membrane recruitment and formation of membrane-disrupting pores (Sun et al., 2012; Wang et al., 2014).

Unfortunately, for many pseudokinases there is no data available on the pathways they regulate and their specific interaction partners. Since pseudokinases are often deregulated in diseases, a better understanding of pseudokinase interactions and their regulatory mechanisms would be desirable also for the evaluation of their potential as targets for the development of new therapeutics (Jacobsen and Murphy, 2017; Ribeiro et al., 2019).

Unc-51-like kinase 4 (ULK4) harbors sequence variations and lacks essential residues in all 5 conserved kinase motifs that are important for catalytic activity (**Figure 1A**). In particular, the substitutions of the VAIK lysine with leucine and the HRD histidine with phenylalanine are rare events in the human kinome. In addition, the lack of the essential ATP binding motifs such as VAIK (VAIL in ULK4) and DFG (NFC in ULK4) suggests that ULK4 has compromised ATP binding. However, strong ATP binding has recently been reported by a comparative binding study using temperature shift assays (Lucet and Murphy, 2017). A recent structure of the catalytic domain in complex with weakly binding inhibitors revealed a canonical kinase fold and may provide the foundation for the development of ULK4 inhibitors that modulate ULK4 scaffolding function (Khamrui et al., 2020).

**Figure 1:**
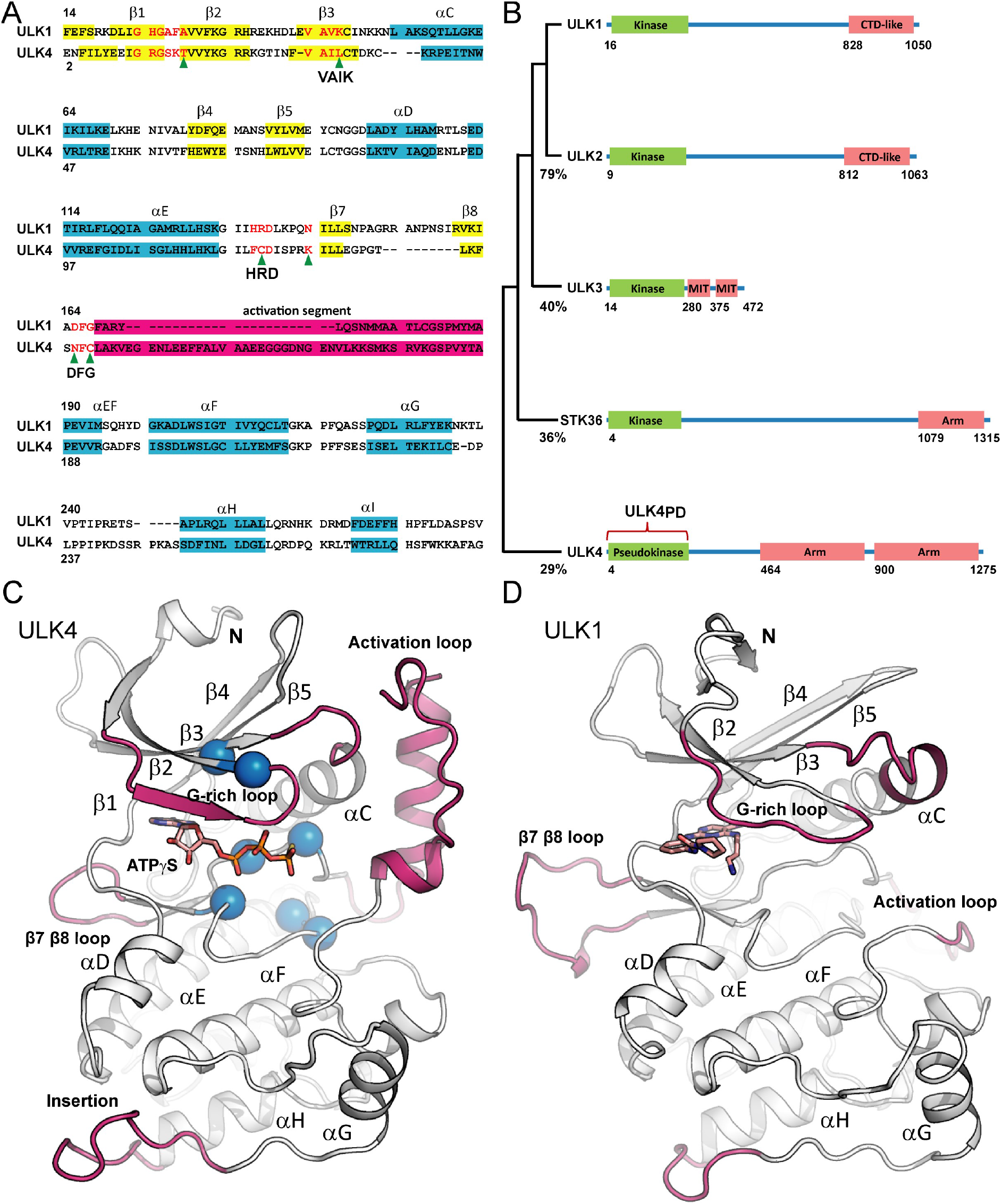
Structural features of ULK kinases. **A:** Structure based alignment of the kinase domains of ULK1 and ULK4. The main secondary structural elements are highlighted and conserved catalytic domain motifs are labelled. Amino acid residue changes in conserved motifs are indicated by a green arrow. **B:** Domain architecture of the human ULK family. Protein interaction domains are labelled as C-terminal domain (CTD; ULK1 and ULK2), microtubule interacting and trafficking molecule (MIT; ULK3) as well as armadillo repeat domain (Arm, STK36 and ULK4). **C:** Structure of the ULK4 pseudokinase domain in complex with ATPγS. Position of residues in conserved catalytic domain motifs that are altered in ULK4 are highlighted by blue spheres. **D:** Structure of ULK1 for comparison. Additional supplemental figures (**Supplemental Figure S2**) show superimpositions of ULK4 with ULK1 and ULK2, respectively.

The ULK family of kinases comprises the catalytically active members ULK1, ULK2, ULK3 and STK36. The kinase domains in ULKs are located at the N-termini of all family members. Usually, the regions C-terminal to the kinase domain contain protein interaction motifs important for substrate recruitment (**Figure 1B**). The role of the C-terminal substrate recruitment domains has been demonstrated for ULK3 (Caballe et al., 2015). Its second microtubule interacting and trafficking (MIT) domain binds to the protein IST1, thus enabling the ULK3 kinase domain to phosphorylate IST1 (Caballe et al., 2015). This mechanism enables ULK3 to regulate the timing of membrane remodeling by the ESCRTIII machinery. Analogously, the ULK1 C-terminal domain-like (CTD-like) domain interacts with autophagy-related protein 13 (ATG13). When activated, the ULK1 kinase domain phosphorylates the bound ATG13, which is part of the cascade ultimately leading to autophagosome formation (Zachari and Ganley, 2017). Despite its lack of kinase activity, a similar recruitment mechanism for substrates or interacting proteins might be also relevant for ULK4 which contains armadillo repeat protein interaction domains. However, no interaction partners for the ULK4 C-terminal armadillo repeats which are also present in C-terminus of STK36 have been thus far reported.

ULK4 has been linked to several human disorders, highlighting the developmental function of this catalytically inactive kinase scaffold protein. Single nucleotide polymorphisms (SNPs) in the ULK4 gene have been linked to blood pressure regulation and hypertension (Levy et al., 2009). As a consequence, certain ULK4 alleles increase the risk of acute aortic dissections (Guo et al., 2016). Genome wide association studies have identified a ULK4 polymorphism associated with increased risk for the development of multiple myeloma (Broderick et al., 2011). Deletion of ULK4 in mice results in hydrocephalus, a condition linked to impaired cilia development (Vogel et al., 2012). Multiple roles for ULK4 in brain development and neuronal function have been reported, including neuronal motility (Lang et al., 2014), cortex development (Lang et al., 2016), hypomyelination (Liu et al., 2018b) and GABAergic signaling (Liu et al., 2018a). However, a molecular mechanism of how ULK4 mediates these cellular signaling functions is lacking. ULK4 falls within a class of “dark” kinases for which there is little or no functional data (Oprea, 2019).

To elucidate the mechanistic basis for ULK4 function, we solved the crystal structure of the ULK4 pseudokinase domain in complex with ATPγS to gain insights into its unique nucleotide binding properties and identified interaction partners using mass spectrometry. The structure of the ULK4 pseudokinase domain revealed a unique nucleotide-binding mode that results in tight metal ion independent binding of the co-factor, also providing a rationale for the disease-relevant polymorphisms affecting ATP binding residues. Through deep evolutionary analysis, we demonstrate that loss of catalytic activity of mammalian ULK4 occurred progressively in evolution with ancestral ULK4 orthologues in plants and protists retaining the canonical active site residues. Remarkably, the selective loss of ULK4 active site residues in mammals and metazoans correlated with the emergence of an extended activation segment that structurally compensates for the loss of the canonical salt bridge between the VAIK lysine and the αC glutamate by stabilizing the regulatory C-helix in an active conformation. Identification of ULK4 binding partners by BioID mass spectrometry suggested roles in microtubule and centrosome function. We propose specific docking/protein-protein interaction sites by delineating the selective constraints imposed on the ULK4 catalytic domain.

## Results and Discussion

Based on secondary structure predictions and sequence alignments with ULK kinases we designed several constructs for expression in *E. coli*. Good yields of soluble protein were observed for the construct containing the pseudokinase domain residues 1-288 (ULK4_PD_). This sequence was expressed in frame with a cleavable (TEV: tobacco etch virus protease) N-terminal purification tag (His_6_-TEV) allowing efficient purification of the recombinant protein. However, removal of the N-terminal tag by TEV protease was difficult, indicating that the cleavage site was not accessible. The tag was therefore not removed for crystallization and functional studies. In size-exclusion chromatography (SEC), ULK4_PD_ eluted as a monomeric protein (**Supplemental Figure S1**). The identity of ULK4_PD_ was confirmed by mass spectrometry, confirming the correct mass of the un-cleaved recombinant protein which however underwent N-terminal methionine excision. No post-translational modifications were detected. The recombinant protein that was more than 95% pure as judged by SDS PAGE readily crystallized in the presence of non-hydrolysable ATPγS yielding crystals that diffracted to 1.9 Å resolution. Data collection statistics and refinement is summarized in **Supplemental Table S1**.

### Structural features of the ULK4 pseudokinase domain

The structure of ULK4_PD_ revealed the canonical bilobal domain architecture of protein kinases (PDB-ID 6TSZ, **Figure 1C**) that was similar to a recent ULK4 structure in complex with an inhibitor (pdb-code: 6U5L) (Khamrui et al., 2020) but a larger portion of the activation segment was visible in the ATP complex. The structures of the active ULK family members ULK1 and ULK2 superimposed well with ULK4_PD_ despite the low sequence identity (29%) shared between these two kinases (Chaikuad et al., 2019; Lazarus et al., 2015) (**Figure 1D, Supplemental Figure S2**). However, major structural differences were observed in the length of the αC helix, the β7-β8 loop length, the length and topology of the activation loop as well as the orientation of the αG helix is also notably different (**Figure 1**). This helix does not form extensive crystal contacts in ULK structures and the ULK4 αG helix remained stable in a molecular dynamics study (see below), suggesting that the captured conformations in the compared crystal structures are also present in solution. The glycine rich loop contains a threonine residue in ULK4 (T16) at a position where in active kinases usually a glycine is found. The introduction of this bulky residue resulted in a rotation of the β1 strand. A similarly rotated β1 strand in combination with a glycine rich loop threonine has been observed in structures of the TYK2, JAK1 and JAK2 pseudokinase domains (Lupardus et al., 2014). The canonical VAIK motif is replaced by VAIL in ULK4 replacing the conserved lysine residue by a leucine (L33). This substitution has not been reported for any other human kinase. In active kinase conformations, the VAIK lysine stabilizes the “in” conformation of helix αC by forming a highly conserved salt bridge with an invariant glutamate present in αC. In addition, this lysine residue is essential for coordinating the nucleotide phosphates in the ATP bound state. Structural superimposition with ULK1 showed that the αC glutamate residue is also missing in ULK4 and it has been replaced by a tryptophan (W46) (**Figure 2A**). The VAIL leucine is however too far away (4 Å) to form efficient hydrophobic interactions with W46. Instead, W46 extends the regulatory spine through interaction with F140, the central residue in the degenerated DFG motif (NFC in ULK4), most likely contributing to the stability of the observed active-like state of the pseudokinase domain. It is interesting to note that the pseudokinase domain of STK40 is the only other human kinase that also bears an NFC motif (Durzynska et al., 2017). Nevertheless, αC is in an active “in” conformation similar to the one observed in phosphorylated ULK1 (pdb code: 4WNO). Helix αC is one turn shorter than in ULK1. However, the length in αC is highly variable in kinases and may depend on the activation state (Eswaran et al., 2007). In ULK2, αC is even shorter than in ULK4 and contains a helical insert in the linker connecting αC with β3 (**Supplemental Figure S2**). The C-terminus of αC is stabilized by interactions of the degenerated NFC motif residue F140 which assumes an NFC “in” conformation.

**Figure 2:**
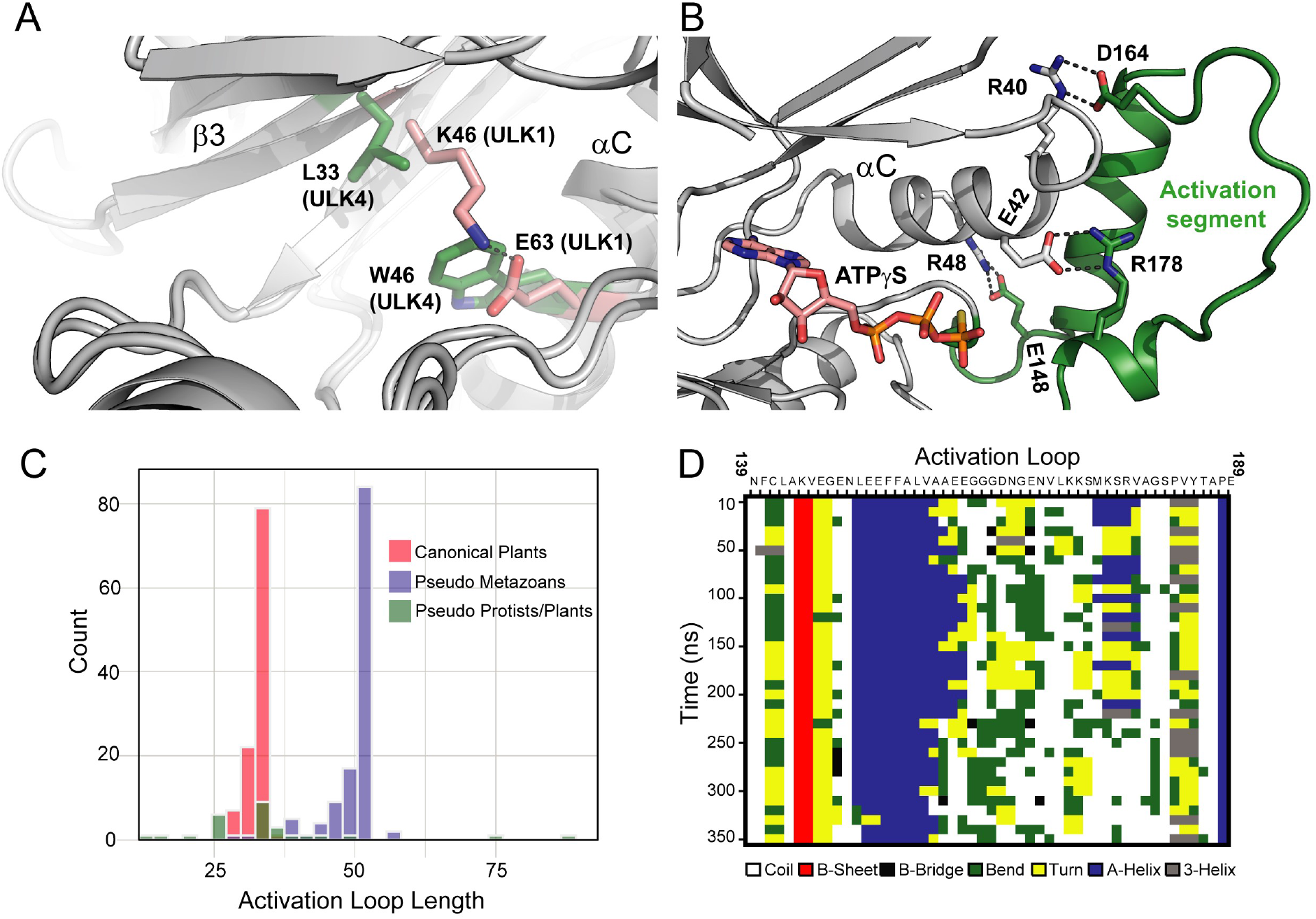
Details of ULK4 structural features and evolutionary conservation. **A:** Comparison of interaction between the ULK1 VAIK motif and αC showing the canonical salt bridge of αC in conformations and ULK4 in which the VAIK lysine is replaced by a leucine residue (L33) and the αC glutamate (W46) by a tryptophan. **B:** Details of the interaction of the ULK4 activation segment helix with αC. Interactions in the catalytic loop are also shown. **C:** Histogram showing the lengths of the activation loop in ULK4 orthologs. Canonical Plants, Pseudo Metazoans, and Pseudo Protists/Plants are colored as light red, light purple, and green, respectively **D:** Secondary structure plot of the activation loop of ULK4. Define Secondary Structure of Proteins (DSSP) was used to define the secondary structure. A maximum likelihood tree of available ULK4 sequences of different species are shown in **Supplemental Figure S3**.

The catalytic loop that comprises the conserved HRD motif in canonical kinases is replaced by an FCD motif in ULK4. However, the phenylalanine (F119) within the FCD motif still anchors the catalytic loop to the kinase core through hydrophobic interactions with residues L109, L112, I122, F137 and F140. The HRD arginine residue that often links the catalytic loop to phosphorylated residues in the activation segment is replaced by a cysteine, but this position is quite variable also in active kinases. The kinase catalytic aspartate, however, is conserved in ULK4. There is no indication that this aspartate can function as a catalytic base, but it still forms a commonly seen salt bridge to the +5 residue (K126) in the catalytic loop. This residue in the very C terminus of the catalytic loop is typically an asparagine in canonical kinases, but it is a lysine (K126) in ULK4. As a result, there is a network of salt bridges linking K126 with the co-factor ATP and D121 in the catalytic loop. To date, the only other pseudokinase for which an aspartate to lysine salt bridge in the catalytic loop has been reported is TRIB1 which maintains a similar catalytic loop conformation as observed in ULK4 (pdb code: 5CEM). The salt bridge is however tilted towards the core of TRIB1 due to a side chain flip of TRIB1 D205 (Murphy et al., 2015).

In comparison with the canonical kinase fold, the ULK4 activation segment contains a large insert with a helical structure (L150 to E161). The activation segment N-terminus contains the unusual degenerated DFG motif (NFC) followed by a typical short sheet region that anchors that activation segment to the lower kinase lobe by hydrogen bonds formed by main chain interactions between two antiparallel short β-sheets. At the N-terminus of the short sheet structure there is a large helix inserted that forms an intricate network of salt bridges with αC possibly stabilizing its inward oriented conformation. The interaction with αC is further stabilized by hydrophobic interactions that form a small hydrophobic core involving residues F153, F154 and L150 at the center of this activation segment helix and P41, L49 located in αC. A salt bride links E148 located in the loop N-terminal to the activation segment helix with αC R48. The hydrophobic environment surrounding this salt bridge suggest a strong polar interaction of the R48/E148 salt bridge (**Figure 2B**). The helix is connected to a coil structure bypassing the catalytic loop. Intriguingly, S182 located in the activation segment 5 residues N-terminal to the APE helix is positioned opposite the catalytic base of the degenerated HRD motif (FCD) mimicking a substrate bound state. Thus, the substrate binding site in ULK4 is blocked by the activation segment. The activation segment terminates with a canonical APE motif.

### Degradation of the active site has co-evolved with an extended activation segment

We performed phylogenetic analysis of ULK4 orthologs from diverse organisms to investigate the origin and evolution of ULK4 as a pseudokinase. We identified ULK4 orthologs from diverse taxonomic groups including metazoans, plants and protists. Phylogenetic analysis of these sequences reveals clear separation of metazoan ULK4 from plants and protists (**Supplemental Figure S3**). Further annotation of the sequences based on the presence or absence of the canonical active residues indicates species specific variations in the canonical active site residues namely K72, E91, D166 and D184 (Protein Kinase A numbering). While metazoan ULK4s lack three of these canonical residues except for the catalytic aspartate (D166) (cyan circular stripes in **Supplemental Figure S3**), orthologues in plant and protists conserve these canonical residues with few exceptions, suggesting that loss of catalytic function of ULK4 emerged later in evolution, presumably for developing specialized metazoan ULK4 pseudokinase functions. In addition to divergence in the canonical active site residues, metazoan ULK4’s displayed variations in the activation loop as well. Comparison of the activation loop lengths across ULK4 orthologues indicates an extended activation segment (mean length of 49.6 amino acids) in pseudo-metazoans compared to ULK4 orthologues from other species (**Figures 2C, S3 and S4**).

The extended segment (L150-E161) forms a helical insert in human ULK4 and packs against the C-helix in the crystal structure. In order to test the stability of this conformation, we performed molecular dynamics (MD) simulation of human ULK4 in the presence of ATP (PDB code: 6TSZ). The simulation indicates that while the inserted helical segment (L150-V157) stably packs against the C-helix, the rest of the loop (A158-E161) is flexible as shown by the lack of secondary structure propensity during the course of simulation (**Figure 2D**). A helical activation loop has also been reported for mouse MLKL (PDB code: 4BTF) (Murphy et al., 2013). This helical segment assumes however a different orientation compared to the ULK4 activation helix. In the mouse MLKL structure, the αC helix is moved outwards making room for the MLKL activation segment helix which resides in a similar position as the ULK4 αC helix. It is interesting to note that in human MLKL, no helical insert has been reported. These data suggest that αC and the activation loop helix may serve pseudokinase specific functions that may even differ between species as observed for MLKL (Petrie et al., 2018). Our evolutionary analysis suggests that the loss of lysine-glutamate salt bridge in metazoan ULK4 required the stabilization of αC by the activation segment helix which in metazoan ULK4 has co-occurred with an extended activation loop, presumably compensating for the loss of the canonical K-E salt bridge. However, since the motif ‘LEEFFALVAAEE’ forming the helical insert in human ULK4 is present in metazoan ULK4 pseudokinases, it is likely that the helical activation loop structure is a common feature of ULK4 sequences that lack the β3-Lysine.

### Evolutionary constraints distinguishing ULK4 from other paralogs

We next performed a Bayesian statistical analysis to identify the evolutionary constraints distinguishing ULK4 from the closely related ULK1-3 and STK36 sequences. This revealed strong ULK4-specific constraints imposed on residues in the N-terminal ATP binding lobe of the pseudo kinase domain. Specifically, these constraints map to residues in the β2-β3 loop as well as in β4 and β5 strands. Some of these constrained residues include R22, R23, and K24 that form a solvent exposed surface patch with hydrophobic residues I27 and F29, which are also ULK4-specific (**Figure 3A**). In addition, an extended network of ULK4 specific residues (F61, W64, E66, and L71) structurally couple the surface patch to the C-helix. These residues are conserved across diverse ULK4 orthologs (including plants and protists) and divergent from the closely related ULK1-3 and STK36 (**Figure 3B**), implying that they are constrained for important ULK4-specific functions. A possible role of this ULK4-specific surface patch could be that it serves as a docking site similar to the one reported for Aurora A with the microtubule associated protein TACC3 (Burgess et al., 2018).

**Figure 3:**
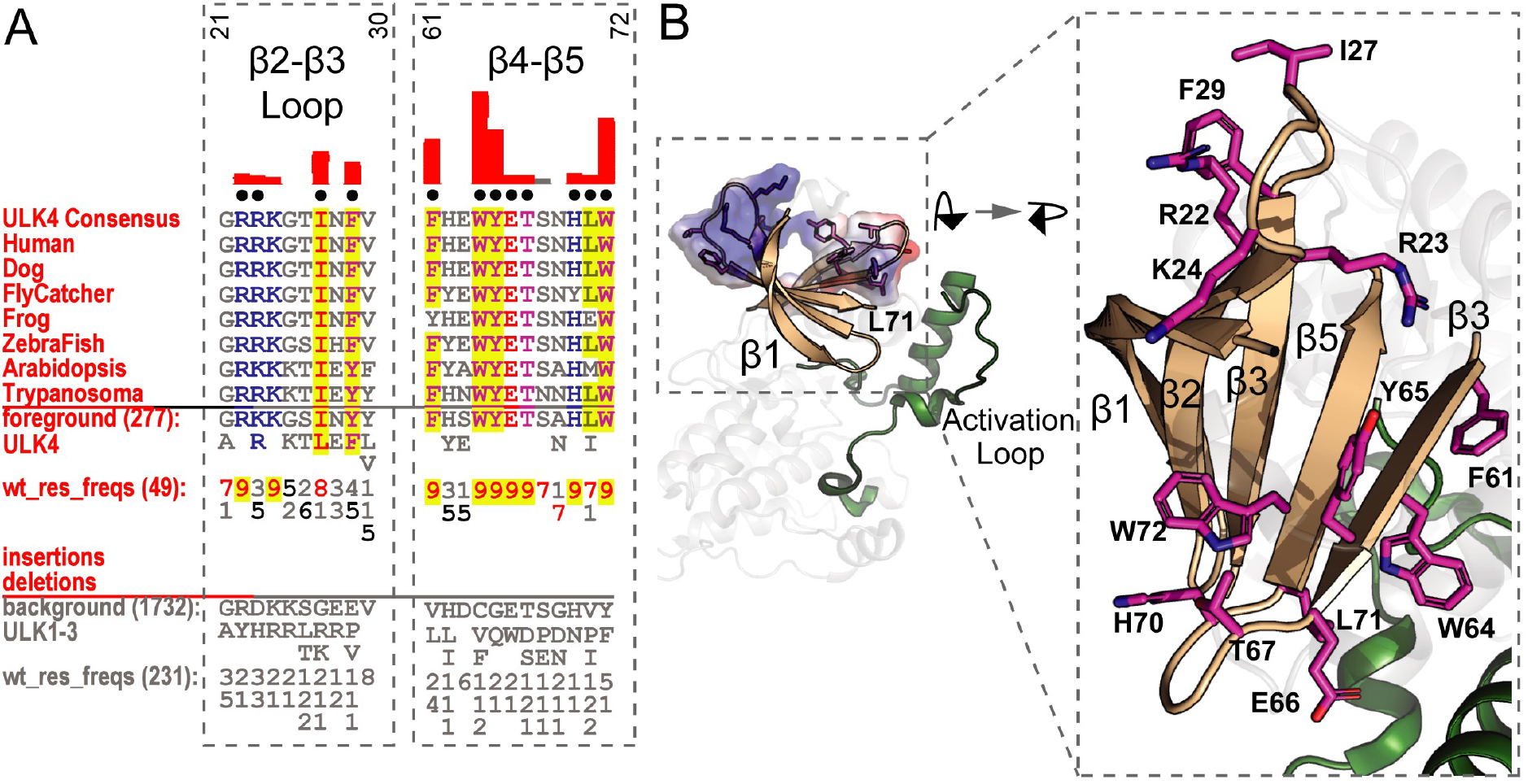
ULK4 specific constraints are in the N-lobe. **A:** Sequence constraints that distinguish ULK4 sequences from closely related ULK1-3 sequences are shown in a contrast hierarchical alignment (CHA). The CHA alignment shows selected ULK4 sequences from diverse organisms as the display alignment. The foreground alignment consists of 277 ULK4 sequences while the background alignment contains 1732 ULK1-3 sequences. The foreground and background alignments are shown as residue frequencies below the display alignment in integer tenths (1–9). The histogram (red) above the display alignment indicates the extent to which distinguishing residues in the foreground alignment diverge from the corresponding position in the background alignment. Black dots mark the alignment positions used by BPPS (Neuwald, 2014) procedure when classifying ULK4 from other ULK sequences. Alignment number is based on the human ULK4 sequence. **B:** ULK4 specific residues mapped onto the crystal structure of human ULK4. The distinguishing residues are shown as sticks and surface electrostatic. The activation loop is colored green. A detailed view is shown in the right panel. Carbon atoms of the residues are colored in magenta. A consensus sequence of ULK4 activation segment is shown in **Supplemental Figure S4**.

### ULK4 has an unusual ATP binding mode

Because conserved motifs known to be essential for nucleotide binding are absent in ULK4, we were particularly interested in how ULK4 interacts with the co-factor ATP. After purification of the recombinant protein the absorbance spectra of ULK4_PD_ showed an unusual ratio at 260 and 280 nm (A_260_/A_280_) suggesting co-purification of nucleotides. Most proteins exhibit maxima at 280 nm due to the presence of tryptophan residues, with A_260_/A_280_ ratio usually ranging from 0.45 to 0.55. In ULK4_PD_, this ratio was 0.76 (**Figure 4A**). An explanation for an aberrant A_260_/A_280_ ratio might be the amino acid composition of the protein. This is not the case in ULK4_PD_ which contains 2.0% tryptophan residues. Another explanation might be the covalent attachment of a co-factor such as ADP-ribose but this scenario can be excluded because any covalent modification would have been detected in our mass spectrometry analysis. We therefore assume that ULK4_PD_ tightly, but non-covalently, bound to a nucleotide metabolite from *E. coli*, and that this interaction was still present at least partially after dialysis and several chromatographic purification steps.

**Figure 4:**
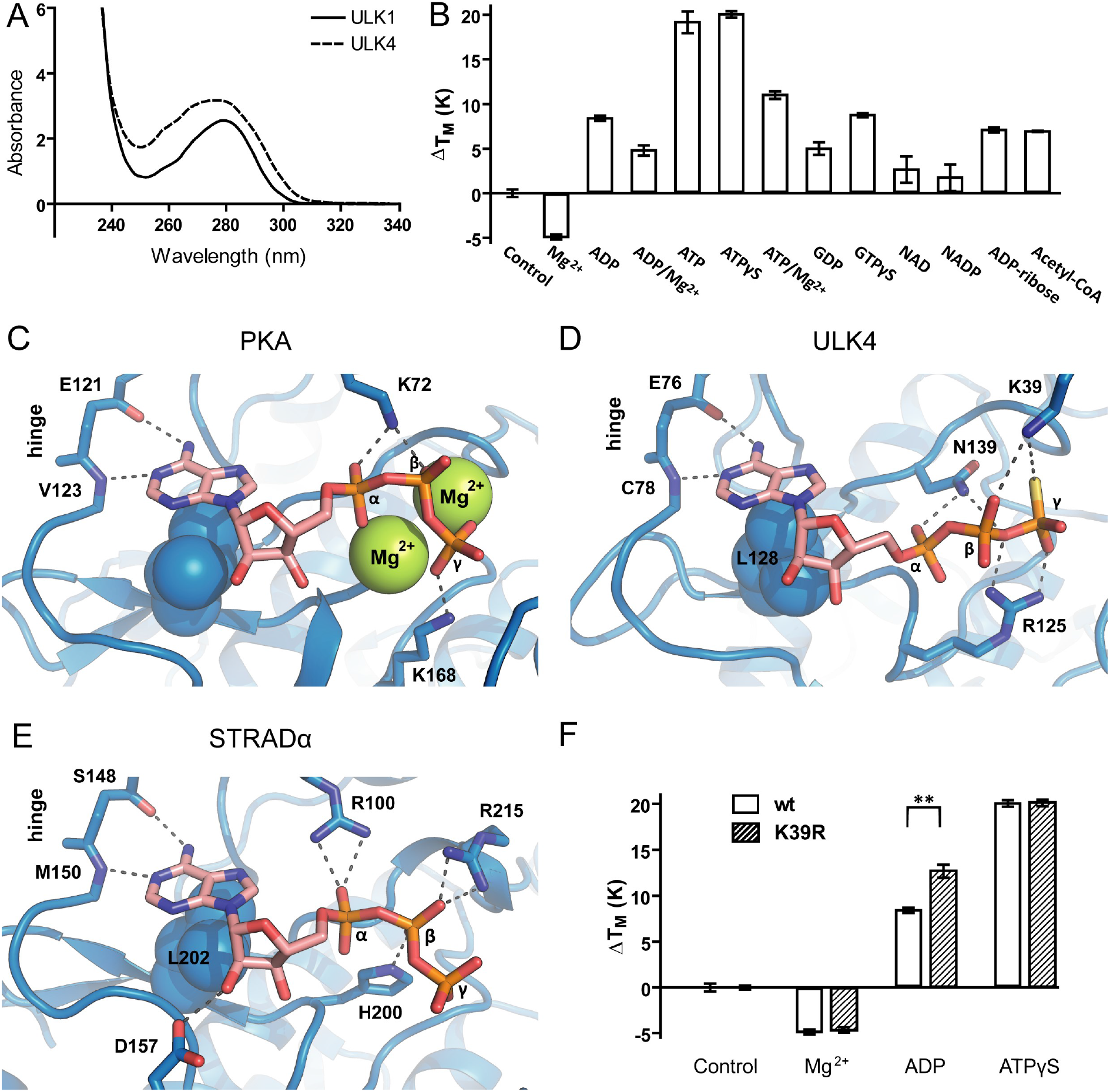
Nucleotide binding to ULK4. **A:** Comparison of UV spectra of ULK1 and ULK4. The high absorbance of ULK4 at 260 nM suggests the presence of ATP after purification from bacteria. **B:** Affinity of nucleotides and effect of metals on binding. Shown are temperature shifts at 1mM nucleotide concentration as well as SEM (n=3). **C:** Canonical binding mode of ATP exemplified by the PKA ATP complex (pdb-ID: 1ATP). Main interactions are shown in stick representation and are labelled. Some side chains have been deleted for clarity. Major hydrophobic contacts interacting with the adenine ring in the low lobe are shown as spheres. **D:** Binding mode of ATPγS in ULK4. Orientations and labels are similar as shown in C. **E:** Nucleotide binding in STRADα (PDB-ID 3GNI). **F:** Effect of K39R polymorphism on nucleotide binding assess by temperature shift experiments. Shown are temperature shifts at 1mM nucleotide concentration as well as SEM (n=3).

In order to assess the durability of ATP binding we performed a Molecular Dynamics (MD) simulation of the crystal structure of ULK4 with ATP bound to it. The co-factor remained stably bound during this 350 ns simulation supporting the high affinity of the co-factor (**Supplemental Figure S5**). A movie illustrating structural changes is also available in the supplemental material.

In order to identify nucleotides that are most likely bound to ULK4 in cells, we performed a co-factor screen using the most common nucleotide-based co-factors and their counter ions. For this analysis we performed thermal shift assays, a screening technology that is well established in our laboratory and that detects molecular interactions based on their effects on protein stability (Fedorov et al., 2012). Recombinant ULK4_PD_ exhibited a melting temperature (T_M_) of 30°C. In comparison to most kinases (40°C<T_M_<60°C) ULK4_PD_ was highly unstable after purification. Both adenine and guanine nucleotides strongly stabilized ULK4_PD_. By far the highest T_M_ shift was observed using ATP (20°C), while the ADP stabilization was 9°C. This highlighted the importance of the ATP γ-phosphate for binding. ADP derivatives such as ADP-ribose and acetyl-CoA stabilized to a similar extent as ADP alone, indicating that the substituents did not interfere strongly with binding (**Figure 4B**). Interestingly, Mg^2+^ ions did not promote nucleotide binding in accordance with previously published results (Murphy et al., 2014). In contrast, the presence of 5 mM Mg^2+^ ions significantly destabilized the ULK4_PD_-ATP complex by 8°C. The pseudokinases calcium/calmodulin-dependent serine protein kinase (CASK), STE20 Related Adaptor Beta (STRADβ), Mixed Lineage Kinase Domain Like Pseudokinase (MLKL) and Ephrin Receptor B6 (EphB6) all lack a functional DFG motif and show metal independent co-factor binding (Bailey et al., 2015; Mukherjee et al., 2008; Murphy et al., 2014). Since ATP is highly abundant in cells, it is likely that the physiological ligand of ULK4_PD_ is indeed ATP. We therefore used the highly stabilizing non-hydrolysable ATP analogue ATPγS for crystallization studies. Screening of a kinase targeted library of 1500 compounds using DSF resulted only in small temperature shifts for some inhibitors (data not shown) suggesting weak interaction with typical kinase inhibitors.

An intriguing aspect of the ULK4_PD_-ATPγS structure was the unprecedented binding mode of the nucleotide. In an active kinase, both the ATP orientation and the triphosphate moiety need to be adjusted perfectly to allow for catalysis. For instance, in the first structure of a kinase ATP complex, protein kinase A (PKA, PDB code: 1ATP), the co-factor binds two Mg^2+^ ions coordinated by the ATP phosphates, the DFG aspartate (D184), the catalytic base in the HRD motif (D166) and N171 and K168 located in the catalytic loop. The glycine rich loop interacts closely with the ATP phosphates forming several main chain interactions. The adenine ring is anchored to the hinge region by two conserved hydrogen bonds with main chain atoms of E121 and V123 (**Figure 4C**). While some interactions between ULK4 and ATPγS are inspired by the kinase ATP binding mode, other interactions have not been described in active kinases and pseudokinase co-factor complexes before (**Figure 4D**). Similar to active kinases, the ATP adenine is anchored to the ULK4 hinge main chain by hydrogen bonding to E76 and C78. Hydrophobic interactions stabilize the adenine ring system which is sandwiched between hydrophobic residues in the β2 strand (V18), the VAIL motif (A31 and L33) as well as the β7 strand (L128). Intriguingly, the lack of the VAIK motif lysine is compensated by K39, capping the αC helix. The ATPγS triphosphate is embedded into an intricate hydrogen-bonding network involving side chains from the tip of αC helix (K39), the catalytic loop (R125 and K126) and the DFG motif (N139). As a result, the ATPγS molecule decorates the back of a groove in ULK4_PD_ with the ribose hydroxyl groups and several phosphate oxygen atoms exposed. A similar groove is observed in the pseudokinase STRADα. Binding of both ATP and the allosteric activator protein MO25 allegedly forces STRADα into an active-like conformation (**Figure 4E**). The ternary complex then binds to and activates the client kinase STK11/LKB1 (Serine/Threonine Kinase 11/Liver Kinase B1) (Zeqiraj et al., 2009b). In this process, the role of ATP is solely to maintain STRADα in its binding-competent form. The ATP molecule interacted directly neither with the client kinase nor with the activator.

Recently, genome-wide association studies (GWAS) linked several single-nucleotide polymorphisms (SNPs) in ULK4 to diseases such as high blood pressure (Ehret et al., 2016; Levy et al., 2009) and sporadic thoracic aortic dissection (STAD) (Guo et al., 2016). Intriguingly, SNP rs2272007 is localized in the pseudokinase domain of ULK4 affecting residue 39. As described above, in the most common ULK4 variant, K39 takes over the role of the missing VAIK lysine. According to “The 1000 Genomes Project”, 68% of sequenced individuals carry the nucleotide T in rs2272007 and thus express ULK4 protein with a lysine in position 39, and 32% carry the nucleotide C and express an arginine in this position. We were therefore interested in how the R39 polymorphism could affect ATP binding, and generated a site-directed mutant carrying this mutation. We also produced another reported variant: N139L. This variant protein was unstable during purification and precipitated quickly, precluding analysis. However, since N139 replaces the aspartate in the degenerated ULK4 DFG motif and the N139 side chain form intricate interactions with the ATP alpha and beta phosphates, the mutation into a hydrophobic lysine is likely to severely compromise ATP binding affecting structural stability of the ULK4 pseudokinase domain. Temperature shift experiments using the R39 variant showed that the un-ligated protein has similar stability as the K39 protein, and the presence of 1 mM ATP resulted in similar temperature shifts suggesting that both lysine and arginine can form efficient salt bridges to the co-factor and that there was enough space in the ATP binding pocket to accommodate the slightly bulkier arginine side chain. Interestingly, a significant difference was observed in ADP binding. While ULK4_PD_ K39 was stabilized by only 9°C, ULK4_PD_ R39 was stabilized by 13°C, suggesting tighter binding of ADP. It is likely that the side chain of K39 is too short to form efficient salt bridges with the ADP β-phosphate, but R39 has sufficient length to form more efficient contacts (**Figure 4F**). Thus, ULK4 K39 and R39 differed in their affinities for ADP whereas the variant N139L was highly unstable due to compromised ATP binding highlighting the importance of ATP binding for ULK4 function and making a compelling case that altered nucleotide affinity is possibly responsible for disease development as suggested by GWAS studies.

### BioID identifies interaction partners of ULK4

To date, no validated ULK4 interacting partners have been reported. Knockdown studies revealed a role in neural cell migration pointing to a cytoskeletal function as well as stimulating role on MAPK signaling (Lang et al., 2014). To better understand the ULK4 interaction landscape, we performed BioID (proximity-dependent biotin identification) (Roux, 2013), a proximity-dependent labelling technique that identifies proteins in the vicinity of a target polypeptide in living cells. The technique makes use of a mutant form of the *E. coli* biotin ligase enzyme BirA* (R118G), which is fused to a protein of interest. The R118G BirA mutant is unable to maintain an interaction with the activated biotinoyl-AMP molecule. This highly reactive intermediate is thus released into the vicinity of the fused bait protein and reacts with amine groups of nearby polypeptides. Biotinylated proteins can then be purified using streptavidin, and identified by mass spectrometry (Gingras et al., 2019). N- and C-terminally FlagBirA* tagged full length ULK4 was expressed in Flp-In T-Rex 293 cells. Immunofluorescence (anti-Flag) indicated that the tagged ULK4 protein is localized in the cytoplasm (**Supplemental Figure S6**), in agreement with data available at the human protein atlas (https://www.proteinatlas.org).

ULK4 BioID was performed using both N- and C-terminally tagged FlagBirA* fusion proteins, and each analysis was performed twice. Data were compared to BioID conducted on 293 Flp-In T-Rex 293 cells expressing the FlagBirA* tag alone to control for endogenously biotinylated proteins and polypeptides that interact non-specifically with the streptavidin-sepharose matrix material. Data were analyzed using the SAINT (statistical analysis of interactomes (Choi et al., 2011; Teo et al., 2014)) algorithm to identify high confidence proximity interactors (displaying a Bayesian false discovery rate ≤0.01 **Figure 5**; **Supplemental Table S2)**. The proximity interactor detected with the highest number of peptides was calmodulin regulated spectrin associated protein 1 (CAMSAP1), a large cytoplasmic scaffolding polypeptide that binds microtubules via its CKK domain, and controls non-centrosomal MT minus-end dynamics (Atherton et al., 2019; Baines et al., 2009). CAMSAP family members are involved in the regulation of a number of key cellular functions, including cell polarity, regulation of neuronal differentiation and axonal regeneration (Akhmanova and Hoogenraad, 2015; Marcette et al., 2014; Pongrakhananon et al., 2018). The related family member CAMSAP3 was also identified as a ULK4 proximity interactor. Intriguingly, many tubulin binding and centrosomal proteins were also detected, including: HAUS2 and HAUS8 (HEC1/NDC80-Interacting Centrosome-Associated Protein), components of the microtubule-binding augmin/HAUS complex, which is essential for mitotic spindle assembly (Goshima et al., 2008; Uehara et al., 2009); centrosomal proteins such as CCP110 (Centriolar Coiled-Coil Protein 110), CEP97 (Centrosomal Protein 97), CSPP1 (Centrosome and Spindle Pole Associated Protein 1) and OFD1 (OFD1 centriole and centriolar satellite protein); and motor proteins of the kinesin family (KIF1B, KIF3B and KIFAP3).

**Figure 5.**
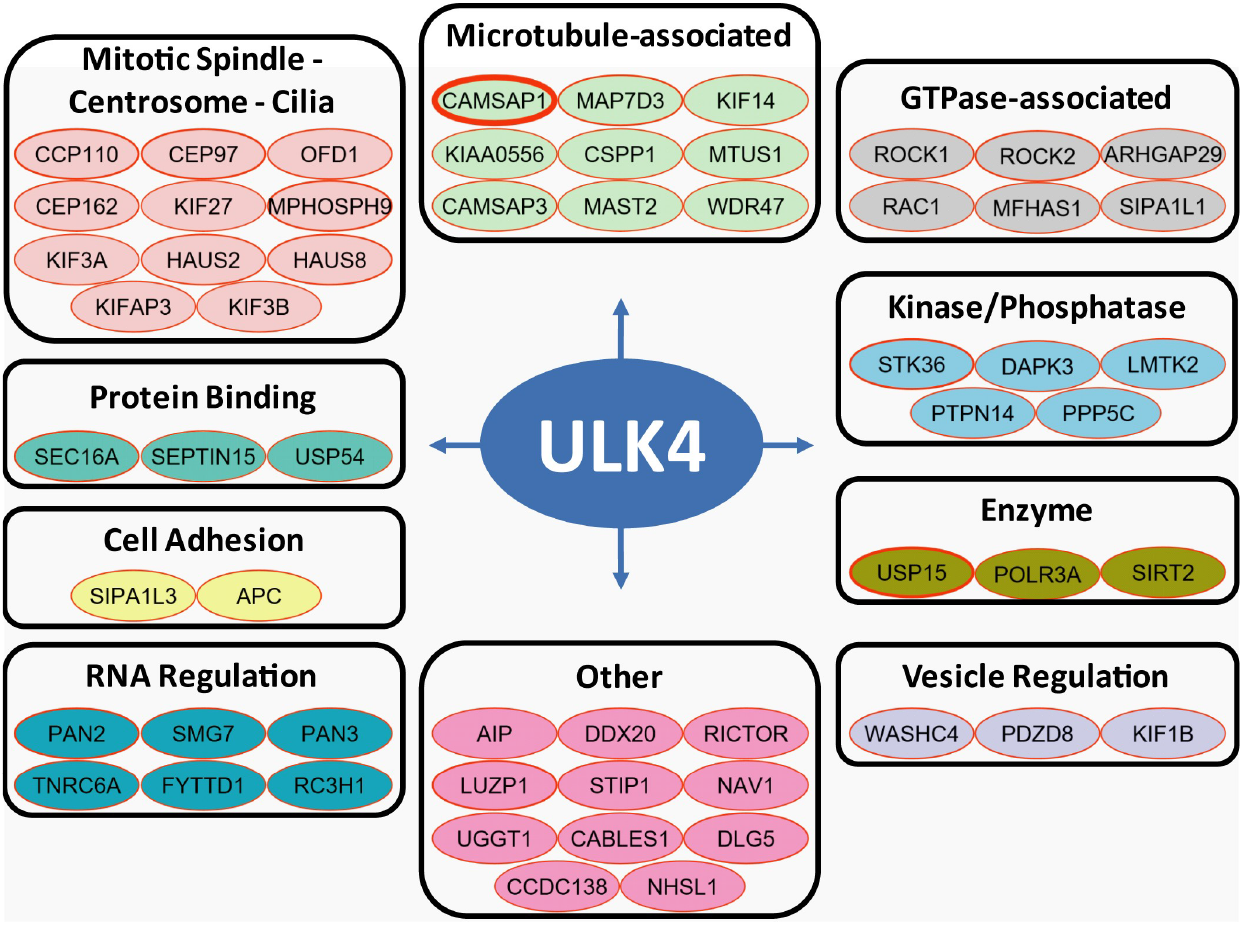
ULK4 Proximity Interaction Network. Proximity interactors of FlagBirA*-ULK4 and/or ULK4-BirA*Flag fusion proteins expressed in Flp-In T-REx 293 cells are shown. Probability of specific interactors was determined using Bayesian statistics against a panel of FlagBirA* only BioID samples. Proteins with a Bayesian FDR of interaction of ≤0.01 were considered statistically significant putative proximity interactors. Border (in red) thickness of nodes is proportional to the number of PSMs for each protein. Proteins are categorized according to GO-based functions. ULK4 cellular localization data are shown in **Supplemental Figure S6**.

Other interacting partners included several protein kinases implicated in microtubular function, such as the Microtubule Associated Serine/Threonine Kinase 2 MAST2, Rho associated coiled-coil containing protein kinase 2 (ROCK2) and Rho GTPase activating protein 29 (ARHGAP29). In addition, PTPN14 (protein tyrosine phosphatase, non-receptor type 14) and the ULK family member STK36, an important regulator of the sonic hedgehog (Shh) pathway (Han et al., 2019; Murone et al., 2000), were also detected as high confidence proximity interactors.

Consistent with the BioID data, CAMSAP1, PTPN14, ROCK1 and ROCK2 co-migrated with full length C-terminal Flag-tagged ULK4 in analytical ultra-centrifugation density gradients, as detected by western blotting (**Figure 6A**). Co-immunoprecipitation using two different methods (C-terminal FLAG-tag IP and streptavidin pulldown) validated the ULK4 interactions with CAMSAP1, ROCK1, ROCK2 and PTPN14 (**Figure 6B and 6C**). The same interactions were validated using N-terminal FLAG-tag constructs with similar results (**Supplemental Figure S7**)

**Figure 6.**
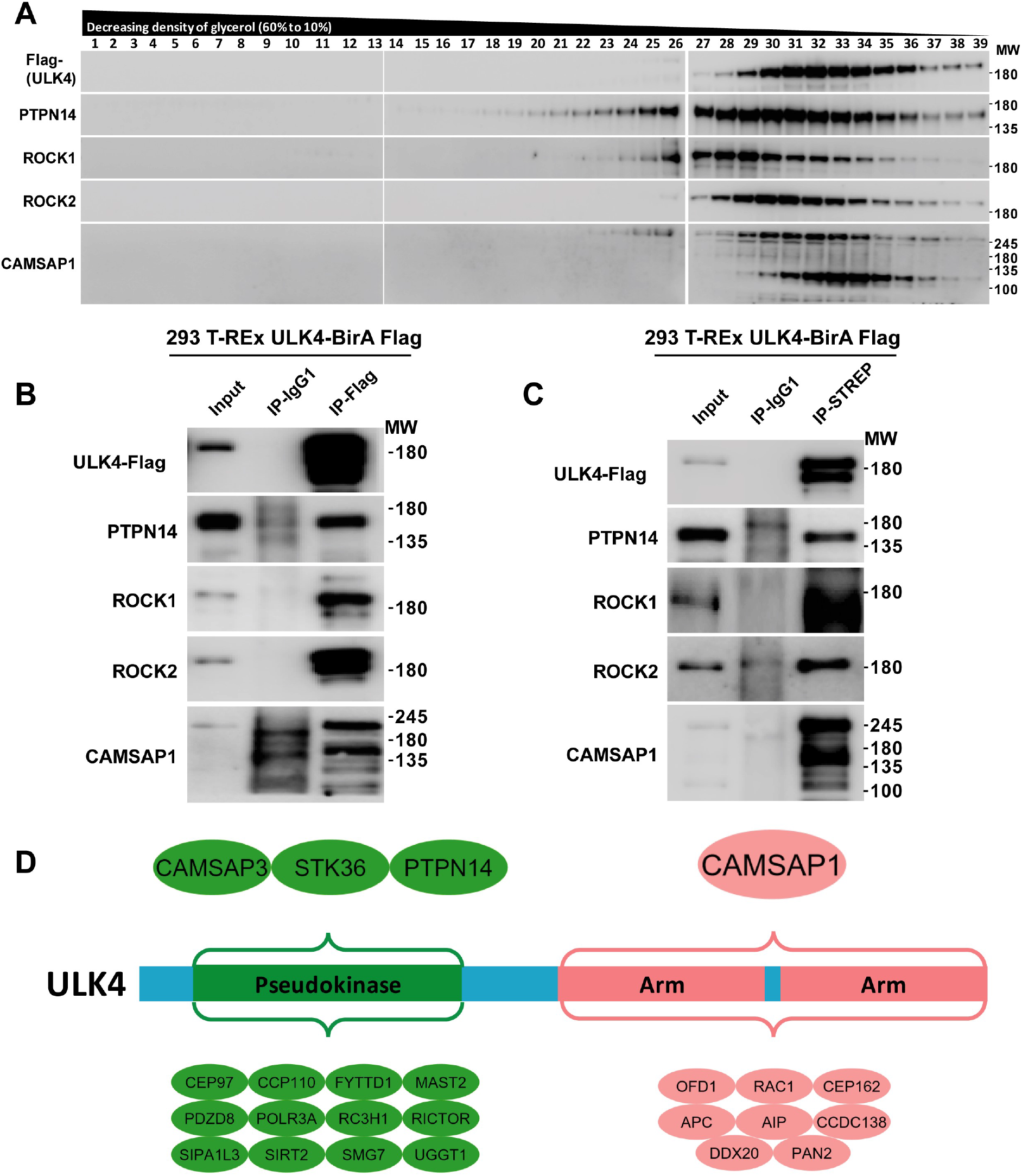
Association studies using density gradients and immunoprecipitation. **A:** Whole cell lysate from tetracycline-treated Flp-In T-REx 293 cells expressing C-terminally FLAG-tagged full-length ULK4 (ULK4-BirA*Flag) was used for density gradient ultra-centrifugation. Immunoblot showed that fractions containing ULK4 also contained PTPN14, ROCK1, ROCK2 and CAMSAP1. **B:** ULK4 was immunoprecipitated from tetracycline-treated Flp-In T-REx 293 cells expressing full-length ULK4-BirA*Flag. Subsequent immunoblotting showed co-immunoprecipitation of PTPN14, ROCK1, ROCK2 and CAMSAP1. **C:** Flp-In T-REx 293 cells expressing tetracycline-inducible Flag- and BirA-tagged full length ULK4 (ULK4-BirA*Flag) were treated with biotin for 24 hrs. Proteins thus biotinylated were pulled down using streptavidin-conjugated beads. Immunoblot showed the presence of PTPN14, ROCK1, ROCK2 and CAMSAP1 in the eluate. **D:** Domain Mapping of ULK4 Proximity Interacting Proteins. Proximity interactors of FlagBirA*-ULK4 and/or ULK4-BirA*Flag pseudokinase domain or Armadillo repeats (Arm) domains fusion proteins expressed in Flp-In T-REx 293 cells are shown. Probability of specific interactors was determined using Bayesian statistics against a panel of FlagBirA* only BioID samples. Proteins with a Bayesian FDR of interaction of ≤0.01 were considered statistically significant putative proximity interactors.

To map the regions of ULK4 that mediate these interactions, the pseudokinase domain and the armadillo repeat region were separately subjected to BioID (**Figure 6D**). This analysis revealed that the armadillo repeat region interacted uniquely with CAMSAP1, OFD1, poly(A) specific ribonuclease subunit 2 (PAN2) and several other proteins. ULK4 pseudokinase domain-specific interactions were detected for STK36, PTPN14, CAMSAP3. The interaction of ULK4 pseudokinase domain with STK36 is particularly intriguing as similar to STRAD/LKB1 this interaction might indicated that ULK4 directly regulates an active kinase (Zeqiraj et al., 2009a). All domain specific detected interactions are summarized in **Supplemental Table S3**. In order to determine whether the polymorphism at position 39 affects the ULK4 protein interaction network that might explain the pathogenicity of this amino acid alteration, the R39 variant was also subjected to BioID. No significant changes in the interactome were detected (**Supplemental Table S4**). Based on our DSF binding study, the R39 variant also bound ATP with similar affinity to the K39 protein. Stability of the pseudokinase domain fold is thus unlikely to be affected by this amino acid change.

The presented structural, bioinformatics, biochemical data together with our BioID interaction study provided a diversity of high confidence domain specific interactions and suggested a microtubular and centrosomal function of ULK4. The high affinity interaction with ATP is required for the stability of the pseudokinase domain, maintaining these interactions. The high-resolution structure of the ULK4 ATPγS complex revealed a highly unusual ATP binding mode despite the lack of canonical ATP interaction motifs demonstrating how this unusual pseudokinase redesigned interaction for co-factor binding. Our evolutionary studies indicated that ULK4 orthologs in plants and protists conserve the canonical active site residues, suggesting that they function as ‘active’ kinases. In contrast, the metazoan-specific variations in the active site suggest that loss of kinase activity and emergence of ULK4 as a pseudokinase occurred later in evolution, presumably to accommodate the specialized functions of differentiated cell types such as the metazoan specific nervous and immune systems. We hope that the presented data will stimulate future research on the function of this poorly studied pseudokinase.

## Significance

ULK4 is an understudied protein associated with diseases such as hypertension and psychiatric disorders. Even though the overall structure of ULK4 has been reported before (Khamrui et al., 2020), we provided in this study important new data comprising structural details of the ULK4 ATP interaction in the absence of conserved canonical ATP binding motifs, evolutionary aspects as well as interaction partners of ULK4. Despite alterations in ATP binding motifs that are essential in canonical kinases, many pseudokinases bind ATP or mimic an ATP bound state which may not require or is even opposed by Mg^2+^ binding (Murphy et al., 2014) (Scheeff et al., 2009). In ULK, polymorphism of key ATP binding residues such as K39R and N139L have been linked to disease development, suggesting that ATP binding is required for ULK4 function. Interaction of the ATP cofactor in canonical kinases leads to a closed structure that is considerably more stable. It is therefore likely that ATP binding and the significant stability increase is required for ULK4 scaffolding function and its role in cellular signaling. Additional structural elements co-emerged with the loss of canonical motifs in evolution that compensate for the loss of stabilizing effects. For instance, our interspecies analysis of ULK4 sequences showed that alteration of the active site motifs in metazoan ULK4 has evolved with an extended activation loop, which stabilizes the flexible C-helix and compensates for the loss of VAIK lysine and the αC glutamate. The diversity of structural mechanisms that show how pseudokinases maintain strong binding activity for ATP in the absence of canonical binding motifs is fascinating and suggest an essential role of stable domain structures in pseudokinases which we confirmed by MD simulations. In order to shed light on signaling pathways regulated by ULK4 we performed interaction studies using BioID mass spectroscopy. These data together with biochemical validation revealed high confidence interaction partners that were mapped to the pseudokinase and to the armadillo repeat domains, respectively. Many of the identified ULK4 interaction partners have centrosomal function suggesting a centrosomal role of ULK4. In addition, a number of active kinases have been identified as ULK4 interactors including the ULK family member STK36. Our studies provide a structural framework for targeting ULK4 in diseases.

## Supporting information

STAR methods & Key Resource Table

Supplemental movie ULK4_ATP_MD

## Acknowledgments

SK, FP, DC, SM are grateful for support by the SGC, a registered charity (number 1097737) that receives funds from AbbVie, Bayer Pharma AG, Boehringer Ingelheim, Canada Foundation for Innovation, Eshelman Institute for Innovation, Genome Canada, Innovative Medicines Initiative (EU/EFPIA EUbOPEN), Janssen, Merck KGaA Darmstadt Germany, MSD, Novartis Pharma AG, Ontario Ministry of Economic Development and Innovation, Pfizer, São Paulo Research Foundation-FAPESP, Takeda, and Wellcome. Funding for NK from the NIH (1U01CA239106-01: A data analytics framework for mining the dark kinome) is acknowledged. SK is grateful for support by the Collaborative Sonderforschungsbereich 1177 Autophagy (SFB1177) at Frankfurt University, as well as the German Cancer Consortium (DKTK). The data collection at SLS has been supported by the funding from the European Union’s Horizon 2020 research and innovation program under grant agreement number 730872, project CALIPSOplus.

## Author Contributions

SK, SM, BR, RR designed research, FP, DC performed protein biochemistry and crystallization, SM solved and refined the ULK4 crystal structure, SS and NK performed the bioinformatics analysis. BR, J-St-G and MS performed the BioID study and validation data on interaction partners. SK, SM and NK wrote manuscript, which was approved by all authors.

