## Supplementary material for "Nucleotide binding, evolutionary insights and interaction partners of the pseudokinase Unc-51-like kinase 4": STAR methods & Key Resource Table

### RESOURCE AVAILABILITY

#### Lead Contact and Material Availability

#### Data and Code Availability

The model and structure factors reported in this study have been deposited in the PDB database under accession code 6TSZ. All raw mass spectrometry data have been deposited in the MassIVE repository (massive.ucsd.edu) with accession ID MSV000084747.

### EXPERIMENTAL MODEL AND SUBJECT MATERIAL

### METHODS DETAILS

#### Cloning

The DNA coding for a $His_6$-tag, a TEV cleavage site and the ULK4 residues 2 to 288 was synthesized (Genscript) and cloned into the expression vector pET-28a, using the NcoI and XhoI restriction sites. From this DNA template, the mutants K39R and N139L were generated by site-directed mutagenesis using the QuikChange kit (Agilent).

#### Protein expression and purification

The expression plasmid was transformed into Rosetta (DE3) competent *E. coli* (Novagen). The expression was performed as previously described (Burgess-Brown et al., 2014). For $ULK4_{PD}$ purification, bacteria were re-suspended in lysis buffer (50 mM HEPES pH 7.4, 500 mM NaCl, 20 mM imidazole, 0.5 mM TCEP, 5% glycerol) and lysed by sonication (35% amplitude, 10 s pulse and 10 s pause during a 20 min pulse sequence). The lysate was cleared by centrifugation and loaded onto a Ni NTA column. After vigorous rinsing with lysis buffer the $His_6$-tagged protein was eluted in lysis buffer containing 300 mM imidazole. Finally, $ULK4_{PD}$ was concentrated and subjected to gel filtration using an AKTA Xpress system combined with an S200 gel filtration column. The elution volume 91.2 mL indicated the protein to be monomeric in solution. The final yield was 10 mg $ULK4_{PD}$/L TB medium.

#### Differential scanning fluorimetry (DSF)

The DSF assay of ULK4 against a set of nucleotides was performed according to a previously established protocol (Niesen et al., 2007). A solution of 2 µM $ULK4_{PD}$ in assay buffer (20 mM HEPES pH 7.4, 150 mM NaCl, 0.5 mM TCEP, 5% glycerol) was mixed 1:1000 with SYPRO Orange (Sigma). The nucleotides to be tested were added to a final concentration of 1 mM. 20 µL of each sample were placed in a 96-well plate and heated gradually from 25°C to 96°C. The fluorescence intensity was monitored using an Mx3005P real-time PCR instrument (Stratagene) with excitation and emission filters set to 465 and 590 nm, respectively. Data was analysed with the MxPro software.

#### Crystallisation of the $ULK4_{PD}$-ATPγS complex

200 nL of a solution containing the protein-ligand complex (12 mg/mL $ULK4_{PD}$, 1 mM ATPγS) were transferred to a 3-well crystallisation plate (SwisSCI), mixed with 100 nL precipitant solution (0.1 M citrate pH 5.9, 15% 2-propanol, 6% PEG4K) and incubated at 4 °C. Crystals appeared after 2 days and did not change appearance after 7 days. They were mounted in

precipitant solution cryoprotected with additional 25% ethylene glycol. Data was collected at Swiss Light Source, analyzed, scaled and merged with Xia2 (Winter, 2010). The structure was solved by molecular replacement with Phaser (McCoy et al., 2005) using a $ULK3_{KD}$ model as a template (PDB ID 6FDY) and refined by iterative model building using the software Coot (Emsley and Cowtan, 2004) and Refmac5 (Murshudov et al., 1997). The model was validated using MolProbity (Chen et al., 2010). The model and the structure factors have been deposited to the protein databank (http://www.rcsb.org/) with the PDB-ID 6TSZ (crystallographic data collection and refinement data are summarized in **Table S1**).

**BioID sample processing**

BioID was carried out as reported previously (Gupta et al., 2015). In brief, lysis buffer (50 mM Tris-HCl pH 7.5, 150 mM NaCl, 1 mM EDTA, 1 mM EGTA, 1% Triton X-100, 0.1% SDS, protease inhibitor cocktail, turbonuclease) was added to frozen cell pellets, incubated with gentle agitation at 4°C for 1 hr, briefly sonicated and centrifuged at 16,000 x *g* for 30 min at 4°C. Supernatants were incubated with 30 µL streptavidin-sepharose beads (GE Healthcare) for 3 h at 4°C with gentle agitation. Beads were washed with $NH_4HCO_3$ (50 mM) prior to overnight digestion with MS-grade, TPCK-treated trypsin (1 µg, Promega) at 37°C. Additional trypsin (0.5 µg) was added, and beads were incubated for 2 hrs at 37°C. Supernatants were collected and beads rinsed with $NH_4HCO_3$ (50 mM). Both fractions were pooled and samples were lyophilized. Samples were reconstituted in HCOOH (0.1%), de-salted on C18 columns and lyophilized.

**Liquid Chromatography – Mass Spectrometry**

Samples were reconstituted in HCOOH (0.1%), loaded on a pre-column (C18 Acclaim PepMap$^{TM}$ 100, 75 µm x 2 cm, 3 µm, 100Å, Thermo Scientific) and separated on an analytical column (C18 Acclaim PepMap$^{TM}$ RSLC, 75 µm x 50 cm, 3mm, 100Å, Thermo Scientific) via high performance liquid chromatography (LC) over a 120-minute, reversed-phase gradient (5-30% $CH_3CN$ in 0.1% HCOOH) running at 250 nl/min on an EASY-nLC1000 pump in-line with a Q-Exactive HF mass spectrometer (Thermo Scientific) operated in positive ion mode ESI. An MS1 ion scan was performed at 60,000 FWHM followed by MS/MS scans (HCD, 15,000 FWHM) of the twenty most intense parent ions (minimum ion count of 1000 for activation). Dynamic exclusion (within 10 ppm) was set for 5 seconds.

**LC-MS Data Processing**

Raw files (.raw) were converted to .mzML format using Proteowizard (v3.0.19311), then searched using X!Tandem (v2013.06.15.1) and Comet (2014.02 rev. 2) against Human RefSeqV104 (containing 36,113 entries). Search parameters specified a parent MS tolerance of 15 ppm and an MS/MS fragment ion tolerance of 0.4 Da, with up to two missed cleavages allowed for trypsin. No fixed modifications were set but deamidation (NQ), oxidation (M), acetylation (protein N-term) and diglycine (K) were set as variable modifications. Search results were processed through the trans-proteomic pipeline (TPP v4.7) and proteins to which ≥2 unique peptides were assigned and an iProphet probability ≥0.9 were considered to be high confidence identifications. For statistical analysis, a Bayesian FDR was assigned to identified proteins using SAINT (v2.5.0; 18 BioID controls compressed to 4). All raw mass spectrometry data have been deposited in the MassIVE repository (massive.ucsd.edu) with accession ID MSV000084747.

**Cloning and generation of cell lines**

The following were cloned into pcDNA5/FRT/TO plasmid using in-fusion HD cloning kit (Clontech): full length ULK4, kinase domain of ULK4, armadillo repeat domain of ULK4, ULK4 with point mutation K39R. Each of the inserts was tagged to Flag and BirA either N-terminally or C-terminally. These plasmids were transfected into Flp-In T-REx 293 cells (Thermo Fisher) using LipoD293 reagent (SignaGen Laboratories), and stable expression cell pools were generated following manufacturer's instructions. Cells were induced using 1 µg/ml tetracycline. For experiments involving biotinylation, biotin was used at a concentration of 50 µM.

**Density gradient ultra-centrifugation**

Flp-In T-REx 293 cells expressing full length ULK4 (upon induction with tetracycline) were lysed in Tris-buffer containing 0.3% CHAPS. Cleared whole cell lysate was carefully layered on top of a continuous glycerol gradient (10%-60%) in Tris buffer. Following ultra-centrifugation at 35000 r.p.m. for 18 hrs at 4°C, fractions were collected, boiled with sample buffer and run on SDS-PAGE for subsequent western blot analysis.

**Antibodies**

Antibodies against Flag tag, PTPN14, ROCK1 and ROCK2 were procured form Cell Signaling Technology, and that against CAMSAP1 were procured form Abcam.

**Immunoprecipitation**

Flp-In T-REx 293 cells expressing full length ULK4 (upon induction with tetracycline) were lysed in Tris-buffer containing 0.3% CHAPS. Whole cell lysate was subjected to pre-clearing, following which it was used for immunoprecipitation using anti-Flag antibody conjugated to agarose beads. For analysis of biotinylated proteins, streptavidin-conjugated beads were used. The eluate in each case was boiled in sample buffer for immunoblotting.

**Immunofluorescence**

Flp-In T-REx 293 cells expressing full length ULK4 (upon induction with tetracycline) was used for staining with anti-Flag antibody. Texas red was used for secondary staining. Confocal imaging was performed with an Olympus IX81 inverted microscope using 60x/1.4 PlanApo oil-immersion objective and FluoView software.

**Phylogenetic tree**

Kinase domains of ULK4 orthologs were aligned with MAFFT V7.450 (Katoh and Standley, 2013). Maximum likelihood tree was generated using FastTree 2.1 (Price et al., 2010). The tree was annotated and visualized using Interactive Tree of Life (ITOL) webserver (Letunic and Bork, 2016).

**Activation loop length distribution**

UniProt sequences of ULK4 orthologs were identified using a previously curated eukaryotic Protein Kinase (ePK) profile (Kannan et al., 2007L; McSkimming et al., 2016; Talevich et al., 2011)using the MAPGAPS tool (Neuwald, 2009). Residues corresponding to the DFG-Asp (N139 in ULK4) and APE-Glu (E189 in ULK4, Uniprot ID: Q96C45) were used as activation loop boundaries. The lengths of the activation loops were then determined for all ULK4 orthologs and the length distribution was plotted using R version 3.6.3 [Link: https://cran.r-project.org/doc/FAQ/R-FAQ.html#Citing-R].

**Molecular Dynamics (MD) of ULK4 with ATP bound**

Unbiased full atom MD simulation of ULK4 was performed on the solved crystal structure using GROMACS 2018.1 software [https://doi.org/10.1016/j.softx.2015.06.001]. Residues with ambiguous electron density were modelled using the Whatif server (Vriend, 1990). Main chain atoms for the missing G163 were modelled using RosettaLoop (Wang et al., 2007) using the cyclic coordinate descent (CCD) protocol. Due to the lack reliable force field parameters for ATPγS, we modelled ATP using ATPγS coordinates as template and performed unrestrained simulations on the ATP bound complex. Both the protein and ATP were parameterized with CHARMM36-March2019 forcefield (Huang and MacKerell, 2013). The protein was solvated with TIP3P water model in a dodecahedron box. In order to neutralize the charge on the protein, sodium and chloride ions were added to the system. Verlet cutoff was used to define neighbour list for non-bonded interactions [doi: 10.1016/j.cpc.2013.06.003]. Particle Mesh Ewald (PME) was used to calculate long-range interactions. Energy minimization was performed with steepest-descent algorithm and then with conjugate descent with Fmax less than 500kJmol-1nm-1. The canonical ensemble was carried out by heating the system from 0 K to 310 K, using velocity rescaling for 100 ps (Bussi et al., 2007). The isothermal–isobaric ensemble (P = 1 bar, T = 310 K) was carried out using the Berendsen barostat for 100 ps [https://doi.org/10.1063/1.448118]. The unrestrained MD productions were collected using a time step of 2 fs after the isothermal–isobaric ensemble. The trajectories were processed and analyzed using the GROMACS built-in tools. Secondary structures for the MD trajectory was defined using DSSP (Allan and Doherty, 1990; Kabsch and Sander, 1983). Structural visualization was performed using PyMOL [PyMOL 2.3.2].

**Identification of ULK4 sequence constraints**

ULK1-4 UniProt sequences were identified and aligned using previously curated hierarchical ePK profiles (See Methods, Activation loop length distribution). Residues distinguishing ULK4 sequences from other ULK paralogs were then identified using the optimal multiple category Bayesian Partitioning with Pattern Selection (omcBPPS) program (Neuwald, 2014).

### KEY RESOURCES TABLE

| REAGENT or RESOURCE | SOURCE | IDENTIFIER |
|---|---|---|
| Antibodies | | |
| Anti-Flag | Cell Signaling Technology | Cat#14793 |
| Anti-PTPN14 | Cell Signaling Technology | Cat#13808 |
| Anti-ROCK1 | Cell Signaling Technology | Cat#4035 |
| Anti-ROCK2 | Cell Signaling Technology | Cat#9029 |
| Anti-CAMSAP1 | Abcam | Cat#86000 |
| Texas Red goat anti-rabbit | ThermoFisher Scientific | Cat#T6391 |
| Bacterial and Virus Strains | | |
| *Escherichia coli* Rosetta | Novagen | Cat#70954 |
| Biological Samples | | |
| none | | |
| Chemicals, Peptides, and Recombinant Proteins | | |
| HEPES | Fisher BioReagents | Cat#BP310-1 |
| NaCl | Fisher BioReagents | Cat#S/3160/65 |
| TCEP | Goldbio | Cat#TCEP25 |
| Imidazole | Alfa Aesar | Cat#A10221 |
| Glycerol | Fisher BioReagents | Cat#G/0650/17 |
| PEG4K | Molecular dimensions | www.moleculardimensions.com |
| ATPγS | Jena Bioscience | Cat#NU-406-5 |
| citrate | Molecular dimensions | www.moleculardimensions.com |

| | | |
|---|---|---|
| 2-propanol | Molecular dimensions | www.moleculardimensions.com |
| Ethylene glycol | Fluka Analytical | Cat#03750 |
| SYPRO orange | Sigma | Cat#S5692 |
| LipoD293 | SignaGen Laboratories | Cat#SL100668 |
| CHAPS | MilliporeSigma | Cat#C3023 |
| tetracycline | MilliporeSigma | Cat#T3383 |
| Biotin | Biobasic | Cat#BB0078 |
| Critical Commercial Assays | | |
| none | | |
| Deposited Data | | |
| ULK4 in complex with ATPgammaS | This paper | PDB: 6TSZ |
| Mass spectrometry data | This paper | ID MSV000084747 |
| Experimental Models: Cell Lines | | |
| Human: Flp-In T-REx 293 cells | Thermo Fisher | Cat#R780-07 |
| Experimental Models: Organisms/Strains | | |
| none | | |
| Oligonucleotides | | |
| Mutagenesis primer pair for ULK4 K39R: GTGCACCGATAAGTGCAGACGTCCGGAGATTACCAACTG CAGTTGGTAATCTCCGGACGTCTGCACTTATCGGTGCAC | Eurofins | https://www.eurofins.com/ |
| Mutagenesis primer pair for ULK4 N139L: GTACCCTGAAGTTCAGCCTCTTTTGCCTGGCGAAAGTG CACTTTCGCCAGGCAAAAGAGGCTGAACTTCAGGGTAC | Eurofins | https://www.eurofins.com/ |
| Recombinant DNA | | |
| pET-28a(+) encoding ULK4 residues 2-288 | Genscript | Synthetic DNA |
| pET-28a(+) encoding ULK4 K39R residues 2-288 | Genscript | Synthetic DNA |
| pcDNA3.1-eGFP encoding fulllength ULK4 | Genscript | CloneID#OHu10418 |
| pcDNA5/FRT/TO encoding full-length ULK4, ULK4 pseudokinase domain, ULK4 armadillo repeat domain, ULK4 K39R | This paper | |
| Software and Algorithms | | |
| MxPro software | Stratagene | https://www.agilent.com/ |
| FluoView software | Olympus | ttps://www.olympus-lifescience.com/ |
| Xia2 | Winter, 2010 | https://www.ccp4.ac.uk/ |
| Phaser | McCoy, 2005 | https://www.ccp4.ac.uk/ |
| Coot | Emsley, 2004 | https://www.ccp4.ac.uk/ |
| Refmac5 | Murshudov, 1997 | https://www.ccp4.ac.uk/ |
| MolProbity | Chen, 2010 | molprobity.biochem.duke.edu |
| Proteowizard (v3.0.19311) | (Chambers et al., 2012) | http://proteowizard.sourceforge.net/ |
| X!Tandem (v2013.06.15.1) | *The Global Proteome Machine Organization* | https://www.thegpm.org/TANDEM/ |

| | | |
|---|---|---|
| Comet (2014.02 rev. 2) | (Eng et al., 2013) | https://sourceforge.net/projects/comet-ms/ |
| Trans-proteomic pipeline (TPP v4.7) | (Deutsch et al., 2015) | http://tools.proteomecenter.org/wiki/index.php?title=Software:TPP |
| SAINT (v2.5.0) | (Choi et al., 2011) | https://omictools.com/saint-tool |
| Gromacs 2018.1 | Berendsen,1995 | http://www.gromacs.org/Downloads |
| MAFFT V7.450 | Katoh, 2013 | https://mafft.cbrc.jp/alignment/software/ |
| R version 3.6.3 | R Core Team, 2013 | https://www.r-project.org/ |
| FastTree 2.1 | Price, 2010 | http://www.microbesonline.org/fasttree/ |
| MAPGAPS 1.0.1 | Neuwald, 2009 | http://mapgaps.igs.umaryland.edu/ |
| OmcBPPS 1.0 | Neuwald, 2014 | http://www.chain.umaryland.edu/omcbpps/ |
| Other | | |
| Mx3005P qPCR system | Stratagene | https://www.agilent.com/ |
| 3 Lens crystallisation plate | SWISSCI | 3W96T-PS |
| In-fusion HD cloning kit | Clontech | Cat#639648 |
| Inverted microscope with 60x/1.4 PlanApo oil-immersion objective | Olympus | IX81 |
| Streptavidin-sepharose beads | GE Healthcare | |
| C18 Acclaim PepMap$^{TM}$ 100 | Thermo Scientific | |
| Q-Exactive HF mass spectrometer | Thermo Scientific | |
