## Supplemental Data for "Nucleotide binding, evolutionary insights and interaction partners of the pseudokinase Unc-51-like kinase 4"

**Content:**


| | |
|---|---|
| Page 2 | Supplemental figure S1: Protein characterization |
| Page 3 | Supplemental figure S2: Structural comparison with ULK1 and ULK2 |
| Page 4 | Supplemental figure S3: Maximum likelihood tree |
| Page 5 | Supplemental figure S4: Sequence comparisons |
| Page 6 | Supplemental figure S5: MD study/ATP binding |
| Page 7 | Supplemental figure S6: ULK4 localization |
| Page 8 | Supplemental figure S7: Association studies |
| Page 9 | Supplemental Table S1: Crystallographic refinement and data collection |
| Page 10 | Supplemental Table S2-4: BioID studies |

#### Supplemental Figure S1

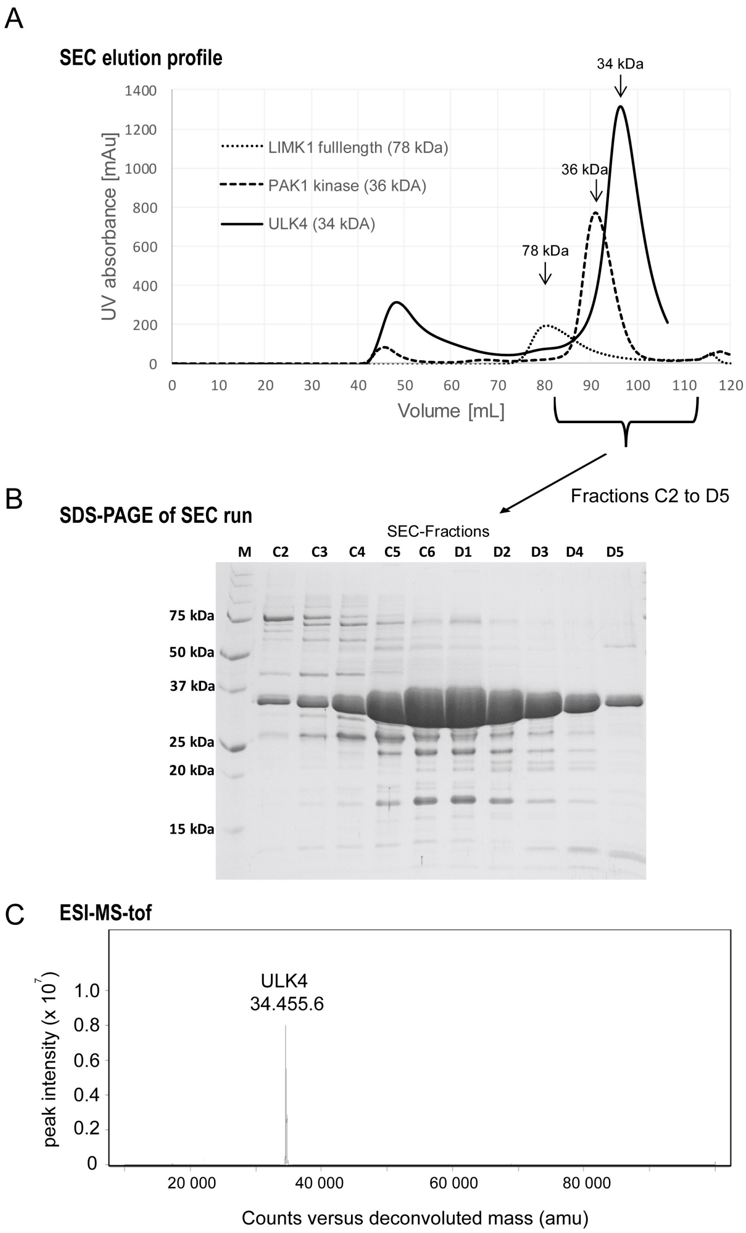


**Supplemental figure S1: Purification of ULK4. A:** Size exclusion chromatography (SEC) of the ULK4 pseudokinase domain and comparison with reference proteins. ULK4 eluted at a volume corresponding to a monomeric protein. **B:** SDS-PAGE of the fractions collected during the SEC run. Molecular weight markers (M) are shown on the right side of the gel image. **C:** Electrospray time of flight mass spectrum (ESI-tof) of purified ULK4. The mass detected corresponded to the expected mass of the recombinant protein including the TEV- His-tag (34585) minus the N-terminal methionine (34455 Da) which was cleaved during expression by N-terminal methionine excision.

#### Supplemental Figure S2

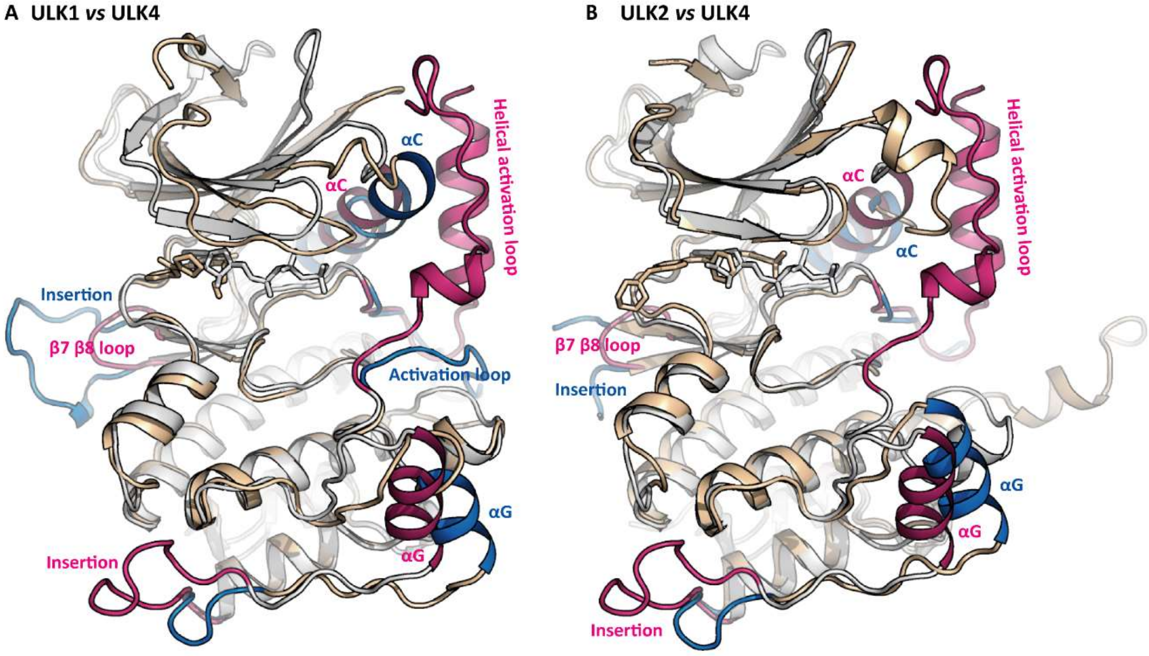


**Supplemental Figure S2: Structural comparison of ULK4 with other ULK family members: A:** Overlay of ULK1 (PDB ID 5CI7, shown in bronze and blue) and ULK4 (6TSZ, grey and red) kinase domains. The fold is well-conserved despite the low sequence identity (29%). Main differences are the helical activation loop in ULK4 and the β7 β8 insertion in ULK1. **B:** Overlay of ULK2 (6QAT, bronze and blue) and ULK4 (6TSZ, grey and red) kinase domains. The ULK2 activation loop is not resolved because it was flexible even in the crystal. In contrast to ULK4, ULK2 contains a short helix bridging the b3 strand and the αC helix.

#### Supplemental Figure S3

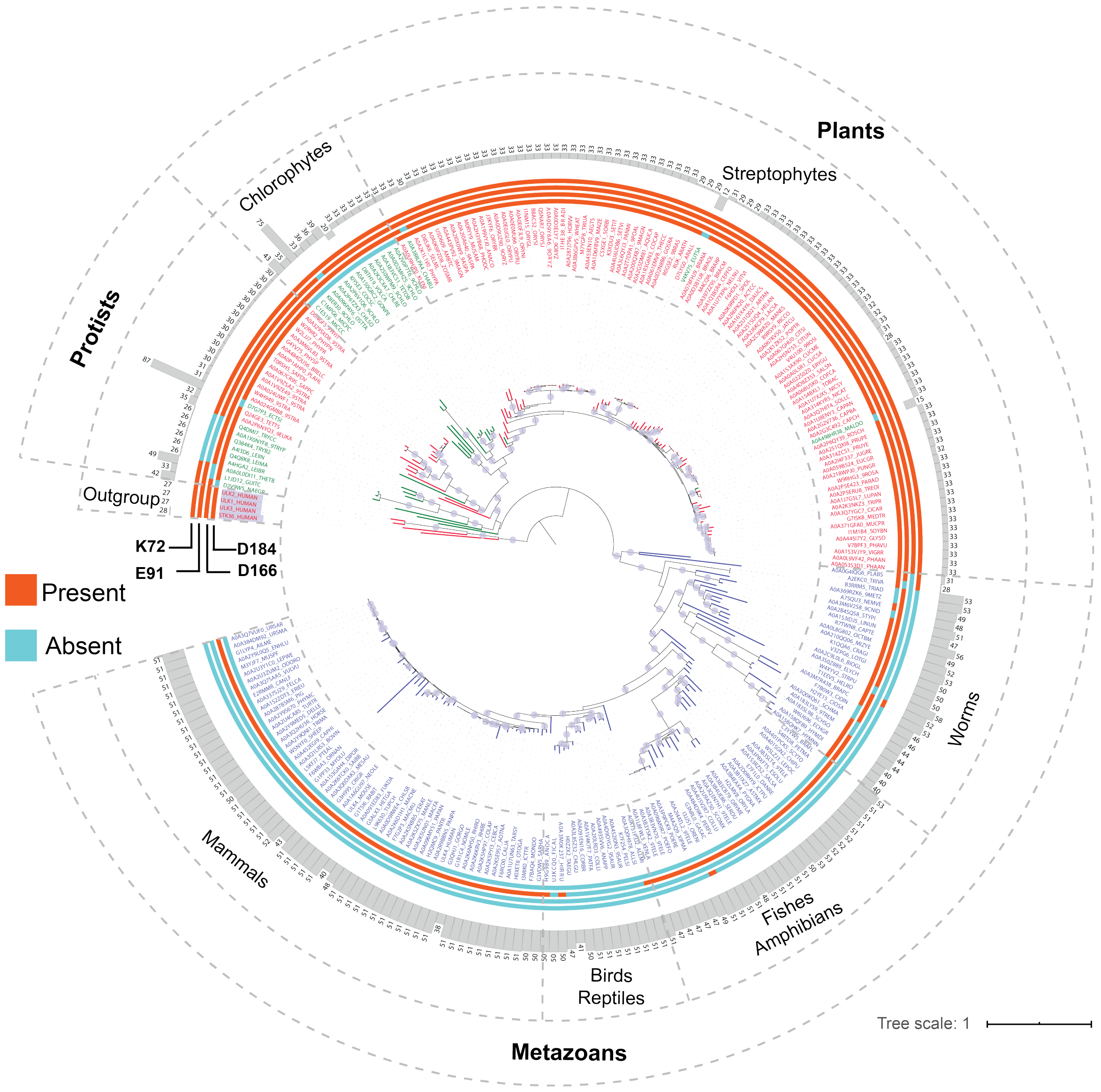


**Supplemental Figure S3: Maximum likelihood tree of ULK4 orthologs**. Presence and absence of key catalytic residues were mapped as stripes (PKA numbering, PDB ID: 1ATP). Activation loop length of the sequences are shown as bar-plot and colored in gray. ULK4 paralogs ULK1-3 and STK36 are used as outgroups. The tree was generated using FastTree 2.1. Nodes with bootstrap values > 0.8 are labelled with circles.

### Supplemental Figure S4

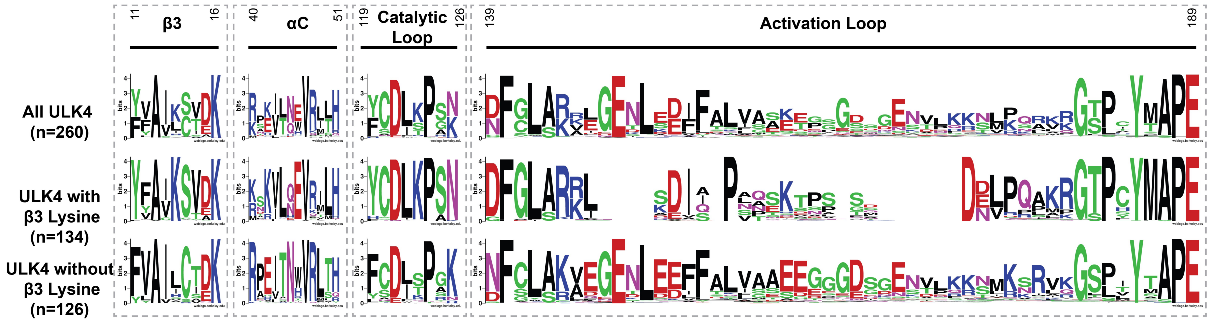


**Supplemental Figure S4: Weblogo of key catalytic motifs in ULK4.** ULK4 orthologs were aligned as in the methods and weblogos were generated using WebLogo Version 2.8 (Crooks et al., 2004). Species specific inserts were removed when generating the weblogos.

#### Supplemental Figure S5

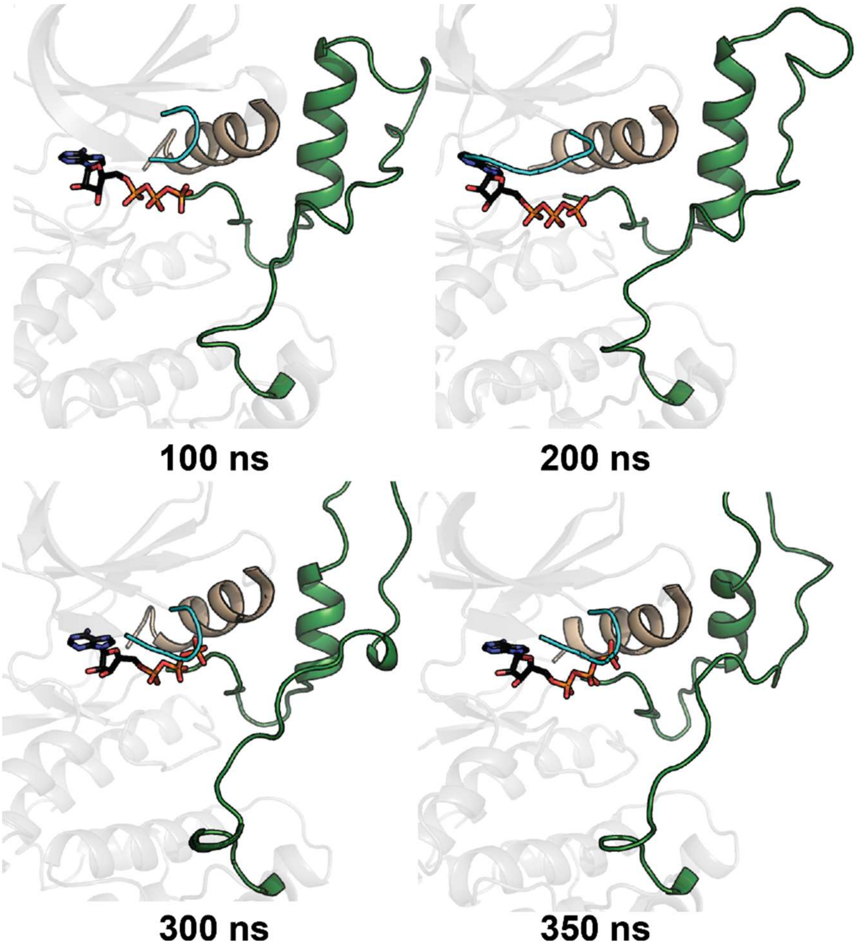


**Supplemental Figure S5. ATP is stably bound to ULK4.** Snapshots of the Molecular Dynamics (MD) simulation of ULK4 crystal structure with ATP bound (PDB ID: 6TSZ). Activation loop is colored forest-green. P-loop and αC-helix are colored in cyan and wheat respectively. ATP is shown as sticks with carbon atoms colored in black.

#### Supplemental figure S6

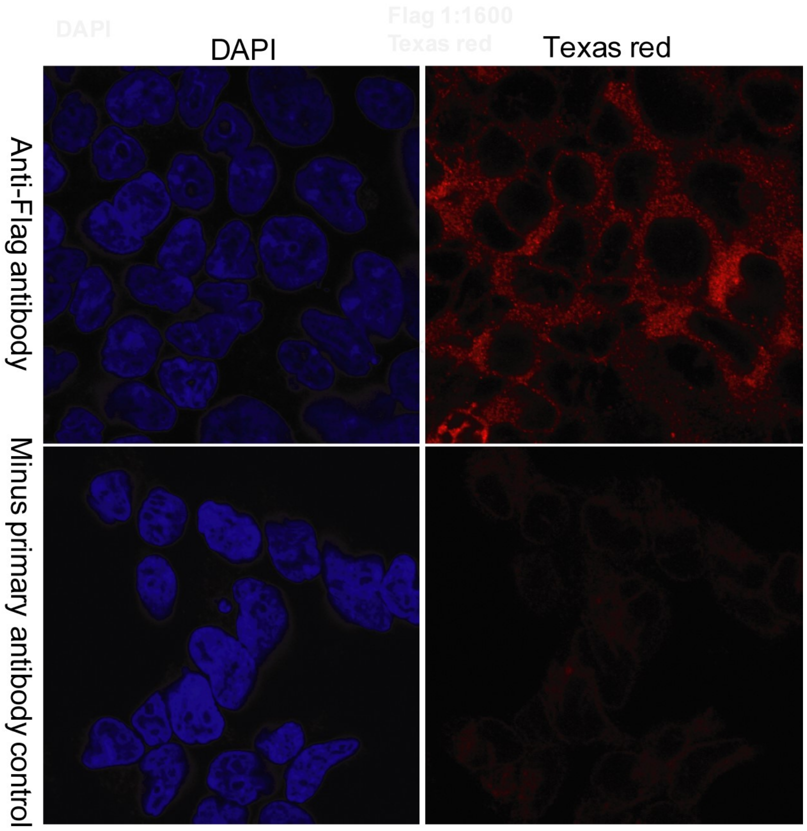


**Supplemental Figure S6: Cellular localization:** 293 T-REx cells expressing tetracycline-inducible C-terminally Flag-tagged full-length ULK4 were stained using anti-Flag antibody. Immunofluorescence showed cytoplasmic localisation of ULK4.

### Supplemental figure S7

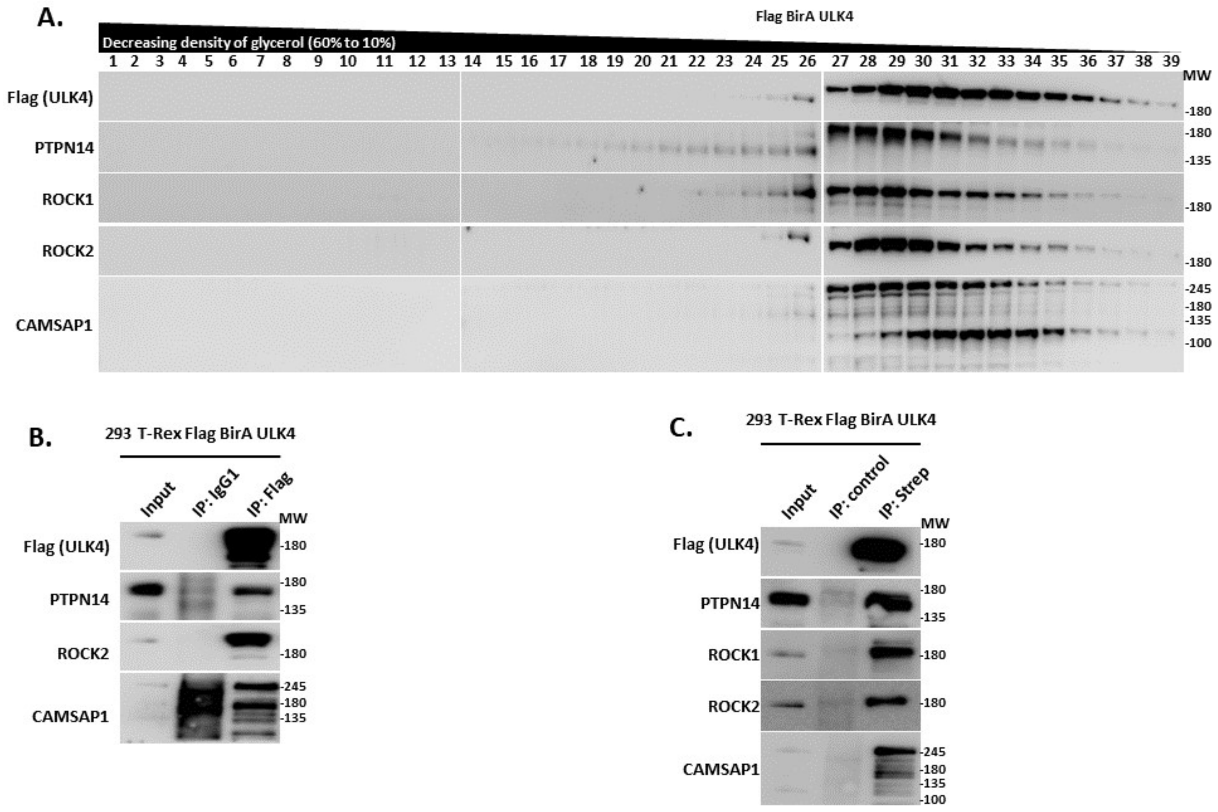


**Supplemental figure S7: Association studies using density gradients and immunoprecipitation. A:** Whole cell lysate from tetracycline-treated Flp-In T-REx 293 cells expressing full-length N-terminal FLAG-BirA tagged ULK4 was used for density gradient ultra-centrifugation. Immunoblot showed that fractions containing ULK4 also contained PTPN14, ROCK1, ROCK2 and CAMSAP1. **B:** Flag-tagged ULK4 was immunoprecipitated from tetracycline-treated Flp-In T-REx 293 cells expressing full-length ULK4. Subsequent immunoblotting showed co-immunoprecipitation of PTPN14, ROCK2 and CAMSAP1. **C:** Flp-In T-REx 293 cells expressing tetracycline-inducible Flag- and BirA- tagged full-length ULK4 were treated with biotin for 24 hrs. Proteins thus biotinylated were pulled down using streptavidin-conjugated beads. Immunoblot showed the presence of PTPN14, ROCK1, ROCK2 and CAMSAP1 in the eluate.

**Supplemental Table S1.** X-ray data collection and refinement statistics.

| | ULK4-ATPγS |
|---|---|
| *Data Collection* | |
| Space Group | *P 1 21 1* |
| *a* (Å) | 62.44 |
| *b* (Å) | 41.47 |
| *c* (Å) | 69.76 |
| $\alpha$ (°) | 90.00 |
| $\beta$ (°) | 113.06 |
| $\gamma$ (°) | 90.00 |
| Molecules/AU | 1 |
| Resolution (Å)[a] | 36.3-1.90 (1.97-1.90) |
| Unique reflections[a] | 25,013 (2518) |
| Completeness (%)[a] | 95.2 (96.4) |
| Multiplicity[a] | 3.4 (3.3) |
| $R_{merge}$ (%)[a] | 6.3 (45.9) |
| CC(1/2)[a] | 0.997 (0.862) |
| Mean $I/\sigma(I)$[a] | 12.7 (2.1) |
| *Refinement* | |
| $R_{work}$, (%)[b] | 20.1 |
| $R_{free}$, (%)[b] | 25.3 |
| No. of atoms | |
| Protein | 2232 |
| Water | 83 |
| Ligands/ions | 31 |
| RMSD bonds (Å) | 0.095 |
| RMSD angles (°) | 1.93 |
| Mean *B* ($Å^2$) | 34.5 |
| PDB entry | 6TSZ |

[a]Values in parentheses are for the highest resolution shell.
[b]$R_{work}$ and $R_{free} = \sum||F_{obs}| - |F_{calc}||/\sum|F_{obs}|$, where $R_{free}$ was calculated with 5% of the reflections chosen at random and not used in the refinement.

#### Supplemental Table S2

| Gene Symbol | Gene Name | controls 1 | 2 | 3 | 4 | SUM | FlagBirA*-ULK4 A1 | A2 | B1 | B2 | SUM | BFDR | ULK4-BirA*Flag A1 | A2 | B1 | B2 | SUM | BFDR |
|---|---|---|---|---|---|---|---|---|---|---|---|---|---|---|---|---|---|---|
| ULK4 | *unc-51 like kinase 4* | 0 | 0 | 0 | 0 | 0 | 4483 | 3785 | 2803 | 2711 | **13782** | na | 2721 | 2528 | 1879 | 1871 | **8999** | na |
| birA | *E. coli biotin carboxylase* | 3726 | 3132 | 3072 | 2815 | 12745 | 4846 | 4041 | 2945 | 2901 | **14733** | na | 2984 | 2642 | 1833 | 1707 | **9166** | na |
| CAMSAP1 | *calmodulin regulated spectrin associated protein 1* | 22 | 18 | 18 | 15 | 73 | 293 | 291 | 289 | 284 | **1157** | **0** | 522 | 515 | 462 | 451 | **1950** | **0** |
| STK36 | *serine/threonine kinase 36* | 0 | 0 | 0 | 0 | 0 | 61 | 52 | 49 | 46 | **208** | **0** | 95 | 94 | 91 | 88 | **368** | **0** |
| LUZP1 | *leucine zipper protein 1* | 19 | 15 | 14 | 14 | 62 | 51 | 46 | 46 | 44 | **187** | **0** | 73 | 69 | 62 | 58 | **262** | **0** |
| HAUS8 | *HAUS augmin like complex subunit 8* | 12 | 11 | 10 | 10 | 43 | 38 | 34 | 33 | 33 | **138** | **0** | 47 | 41 | 41 | 36 | **165** | **0** |
| CCP110 | *centriolar coiled-coil protein 110* | 2 | 2 | 0 | 0 | 4 | 36 | 35 | 35 | 31 | **137** | **0** | 99 | 90 | 87 | 80 | **356** | **0** |
| CEP97 | *centrosomal protein 97* | 5 | 4 | 4 | 3 | 16 | 33 | 32 | 31 | 31 | **127** | **0** | 94 | 93 | 92 | 92 | **371** | **0** |
| MPHOSPH9 | *M-phase phosphoprotein 9* | 2 | 0 | 0 | 0 | 2 | 32 | 30 | 29 | 24 | **115** | **0** | 77 | 68 | 61 | 55 | **261** | **0** |
| PAN2 | *poly(A) specific ribonuclease subunit PAN2* | 6 | 5 | 0 | 0 | 11 | 30 | 28 | 27 | 26 | **111** | **0** | 86 | 80 | 74 | 68 | **308** | **0** |
| ROCK2 | *Rho associated coiled-coil containing protein kinase 2* | 3 | 2 | 2 | 2 | 9 | 27 | 26 | 24 | 24 | **101** | **0** | 68 | 59 | 55 | 52 | **234** | **0** |
| HAUS2 | *HAUS augmin like complex subunit 2* | 8 | 5 | 5 | 5 | 23 | 24 | 24 | 23 | 23 | **94** | **0** | 31 | 30 | 25 | 24 | **110** | **0** |
| SMG7 | *SMG7 nonsense mediated mRNA decay factor* | 6 | 4 | 4 | 4 | 18 | 23 | 22 | 21 | 21 | **87** | **0** | 32 | 28 | 28 | 25 | **113** | **0** |
| UGGT1 | *UDP-glucose glycoprotein glucosyltransferase 1* | 0 | 0 | 0 | 0 | 0 | 21 | 21 | 20 | 15 | **77** | **0** | 7 | 6 | 5 | 4 | **22** | **0** |
| DLG5 | *discs large MAGUK scaffold protein 5* | 2 | 0 | 0 | 0 | 2 | 21 | 20 | 18 | 17 | **76** | **0** | 37 | 30 | 29 | 28 | **124** | **0** |
| TNRC6A | *trinucleotide repeat containing adaptor 6A* | 3 | 3 | 3 | 2 | 11 | 19 | 17 | 16 | 16 | **68** | **0** | 26 | 24 | 23 | 21 | **94** | **0** |
| PTPN14 | *protein tyrosine phosphatase non-receptor type 14* | 0 | 0 | 0 | 0 | 0 | 18 | 17 | 15 | 13 | **63** | **0** | 7 | 6 | 6 | 6 | **25** | **0** |
| KIAA1009 | *centrosomal protein 162* | 0 | 0 | 0 | 0 | 0 | 12 | 11 | 10 | 9 | **42** | **0** | 38 | 37 | 32 | 31 | **138** | **0** |
| SIPA1L3 | *signal induced proliferation associated 1 like 3* | 0 | 0 | 0 | 0 | 0 | 6 | 6 | 5 | 4 | **21** | **0** | 10 | 6 | 6 | 6 | **28** | **0** |
| DAPK3 | *death associated protein kinase 3* | 0 | 0 | 0 | 0 | 0 | 9 | 4 | 4 | 4 | **21** | **0** | 11 | 7 | 5 | 5 | **28** | **0** |
| ARHGAP29 | *Rho GTPase activating protein 29* | 0 | 0 | 0 | 0 | 0 | 6 | 6 | 5 | 4 | **21** | **0** | 7 | 6 | 5 | 5 | **23** | **0** |
| RC3H1 | *ring finger and CCCH-type domains 1* | 0 | 0 | 0 | 0 | 0 | 5 | 5 | 5 | 5 | **20** | **0** | 7 | 4 | 4 | 4 | **19** | **0** |
| PAN3 | *poly(A) specific ribonuclease subunit PAN3* | 0 | 0 | 0 | 0 | 0 | 7 | 4 | 4 | 4 | **19** | **0** | 15 | 14 | 14 | 11 | **54** | **0** |
| SEC16A | *SEC16 homolog A, endoplasmic reticulum export factor* | 10 | 9 | 8 | 7 | 34 | 29 | 29 | 27 | 24 | 109 | 0.02 | 43 | 43 | 40 | 32 | **158** | **0** |
| WDR47 | *WD repeat domain 47* | 0 | 0 | 0 | 0 | 0 | 4 | 3 | 2 | 2 | 11 | 0.03 | 8 | 6 | 5 | 5 | **24** | **0** |
| KIF1B | *kinesin family member 1B* | 2 | 0 | 0 | 0 | 2 | 8 | 8 | 6 | 4 | 26 | 0.05 | 18 | 15 | 14 | 11 | **58** | **0** |
| KIFAP3 | *kinesin associated protein 3* | 0 | 0 | 0 | 0 | 0 | 6 | 6 | 2 | 0 | 14 | 0.08 | 16 | 14 | 14 | 12 | **56** | **0** |
| CSPP1 | *centrosome and spindle pole associated protein 1* | 0 | 0 | 0 | 0 | 0 | 4 | 3 | 2 | 0 | 9 | 0.09 | 17 | 16 | 14 | 13 | **60** | **0** |
| USP54 | *ubiquitin specific peptidase 54* | 0 | 0 | 0 | 0 | 0 | 3 | 3 | 2 | 0 | 8 | 0.09 | 8 | 7 | 6 | 4 | **25** | **0** |
| MAST2 | *microtubule associated serine/threonine kinase 2* | 0 | 0 | 0 | 0 | 0 | 3 | 2 | 2 | 0 | 7 | 0.1 | 7 | 5 | 4 | 3 | **19** | **0** |
| MAP7D3 | *MAP7 domain containing 3* | 14 | 14 | 13 | 13 | 54 | 41 | 39 | 36 | 35 | 151 | 0.12 | 55 | 50 | 50 | 45 | **200** | **0** |
| SIPA1L1 | *signal induced proliferation associated 1 like 1* | 0 | 0 | 0 | 0 | 0 | 2 | 2 | 2 | 0 | 6 | 0.13 | 5 | 5 | 5 | 3 | **18** | **0** |
| KIF3B | *kinesin family member 3B* | 0 | 0 | 0 | 0 | 0 | 3 | 2 | 0 | 0 | 5 | 0.21 | 6 | 6 | 5 | 4 | **21** | **0** |
| OFD1 | *OFD1 centriole and centriolar satellite protein* | 11 | 9 | 6 | 5 | 31 | 23 | 23 | 20 | 19 | 85 | 0.25 | 36 | 32 | 31 | 31 | **130** | **0** |
| MFHAS1 | *malignant fibrous histiocytoma amplified sequence 1* | 0 | 0 | 0 | 0 | 0 | 2 | 2 | 0 | 0 | 4 | 0.25 | 10 | 10 | 8 | 6 | **34** | **0** |
| MTUS1 | *microtubule associated scaffold protein 1* | 8 | 4 | 3 | 3 | 18 | 15 | 11 | 11 | 11 | 48 | 0.41 | 31 | 29 | 26 | 26 | **112** | **0** |
| DDX20 | *DEAD-box helicase 20* | 5 | 4 | 3 | 3 | 15 | 13 | 9 | 9 | 8 | 39 | 0.41 | 19 | 18 | 17 | 17 | **71** | **0** |
| ROCK1 | *Rho associated coiled-coil containing protein kinase 1* | 6 | 0 | 0 | 0 | 6 | 6 | 6 | 6 | 6 | 24 | 0.41 | 42 | 34 | 31 | 31 | **138** | **0** |
| KIF27 | *kinesin family member 27* | 0 | 0 | 0 | 0 | 0 | 2 | 0 | 0 | 0 | 2 | 0.42 | 7 | 7 | 6 | 5 | **25** | **0** |
| NAV1 | *neuron navigator 1* | 4 | 3 | 3 | 2 | 12 | 5 | 5 | 3 | 2 | 15 | 0.71 | 19 | 16 | 14 | 13 | **62** | **0** |
| USP15 | *ubiquitin specific peptidase 15* | 99 | 83 | 79 | 76 | 337 | 50 | 50 | 49 | 45 | 194 | 0.93 | 276 | 248 | 243 | 241 | **1008** | **0** |
| LMTK2 | *lemur tyrosine kinase 2* | 0 | 0 | 0 | 0 | 0 | 0 | 0 | 0 | 0 | 0 | na | 11 | 10 | 9 | 8 | **38** | **0** |
| KIAA0556 | *KIAA0556* | 0 | 0 | 0 | 0 | 0 | 0 | 0 | 0 | 0 | 0 | na | 6 | 5 | 5 | 4 | **20** | **0** |
| CAMSAP3 | *calmodulin regulated spectrin associated protein family member 3* | 3 | 0 | 0 | 0 | 3 | 18 | 15 | 14 | 11 | **58** | **0** | 17 | 14 | 11 | 9 | **51** | **0.01** |
| APC | *APC regulator of WNT signaling pathway* | 3 | 0 | 0 | 0 | 3 | 9 | 8 | 8 | 7 | 32 | 0.07 | 17 | 14 | 12 | 11 | **54** | **0.01** |
| NHSL1 | *NHS like 1* | 2 | 0 | 0 | 0 | 2 | 5 | 4 | 3 | 2 | 14 | 0.28 | 9 | 9 | 9 | 6 | **33** | **0.01** |
| KIF3A | *kinesin family member 3A* | 2 | 0 | 0 | 0 | 2 | 3 | 3 | 3 | 2 | 11 | 0.41 | 11 | 10 | 8 | 6 | **35** | **0.01** |
| KIF14 | *kinesin family member 14* | 10 | 5 | 5 | 4 | 24 | 16 | 14 | 14 | 13 | 57 | 0.6 | 24 | 20 | 20 | 19 | **83** | **0.01** |
| KIAA1033 | *WASH complex subunit 4* | 13 | 6 | 5 | 4 | 28 | 16 | 15 | 14 | 13 | 58 | 0.72 | 32 | 28 | 28 | 27 | **115** | **0.01** |
| CCDC138 | *coiled-coil domain containing 138* | 0 | 0 | 0 | 0 | 0 | 0 | 0 | 0 | 0 | 0 | na | 3 | 3 | 3 | 3 | **12** | **0.01** |
| POLR3A | *RNA polymerase III subunit A* | 0 | 0 | 0 | 0 | 0 | 6 | 3 | 3 | 3 | **15** | **0.01** | 2 | 2 | 2 | 0 | 6 | 0.13 |
| RICTOR | *RPTOR independent companion of MTOR complex 2* | 0 | 0 | 0 | 0 | 0 | 5 | 4 | 3 | 2 | **14** | **0.01** | 2 | 2 | 0 | 0 | 4 | 0.25 |
| RAC1 | *Rac family small GTPase 1* | 0 | 0 | 0 | 0 | 0 | 4 | 3 | 3 | 3 | **13** | **0.01** | 2 | 2 | 0 | 0 | 4 | 0.25 |
| CABLES1 | *Cdk5 and Abl enzyme substrate 1* | 2 | 0 | 0 | 0 | 2 | 10 | 9 | 8 | 8 | **35** | **0.01** | 5 | 4 | 2 | 2 | 13 | 0.29 |
| FYTTD1 | *forty-two-three domain containing 1* | 0 | 0 | 0 | 0 | 0 | 4 | 4 | 4 | 4 | **16** | **0** | 2 | 0 | 0 | 0 | 2 | 0.42 |
| SEPT15 | *Septin15* | 0 | 0 | 0 | 0 | 0 | 4 | 4 | 3 | 3 | **14** | **0** | 2 | 0 | 0 | 0 | 2 | 0.42 |
| PDZD8 | *PDZ domain containing 8* | 4 | 4 | 3 | 3 | 14 | 30 | 28 | 27 | 26 | **111** | **0** | 10 | 7 | 7 | 6 | 30 | 0.59 |
| STIP1 | *stress induced phosphoprotein 1* | 17 | 12 | 12 | 12 | 53 | 44 | 44 | 39 | 38 | **165** | **0** | 7 | 7 | 6 | 6 | 26 | 0.93 |
| SIRT2 | *sirtuin 2* | 0 | 0 | 0 | 0 | 0 | 27 | 24 | 23 | 23 | **97** | **0** | 0 | 0 | 0 | 0 | 0 | na |
| PPP5C | *protein phosphatase 5 catalytic subunit* | 0 | 0 | 0 | 0 | 0 | 4 | 4 | 4 | 3 | **15** | **0** | 0 | 0 | 0 | 0 | 0 | na |
| AIP | *aryl hydrocarbon receptor interacting protein* | 2 | 0 | 0 | 0 | 2 | 8 | 8 | 7 | 7 | **30** | **0.01** | 0 | 0 | 0 | 0 | 0 | na |

**Supplemental Table S2: BioID high confidence proximity interactors for full-length ULK4**

#### Supplemental Table S3

| | | FlagBirA*-ULK4 | | | | | | | | | | | | ULK4-BirA*Flag | | | | | | | | | | | |
|---|---|---|---|---|---|---|---|---|---|---|---|---|---|---|---|---|---|---|---|---|---|---|---|---|---|
| | | K39 | | | | | | R39 | | | | | | K39 | | | | | | R39 | | | | | |
| Gene Symbol | Gene Name | A1 | A2 | B1 | B2 | SUM | BFDR | A1 | A2 | B1 | B2 | SUM | BFDR | A1 | A2 | B1 | B2 | SUM | BFDR | A1 | A2 | B1 | B2 | SUM | BFDR |
| ULK4 | *unc-51 like kinase 4* | #### | #### | #### | #### | ##### | na | #### | #### | #### | #### | ##### | na | #### | #### | #### | #### | #### | na | #### | #### | #### | #### | ##### | na |
| birA | *E. coli biotin carboxylase* | #### | #### | #### | #### | ##### | na | #### | #### | #### | #### | ##### | na | #### | #### | #### | #### | #### | na | #### | #### | #### | #### | ##### | na |
| CAMSAP1 | *calmodulin regulated spectrin associated protein 1* | 293 | 291 | 289 | 284 | **1157** | **0** | 260 | 256 | 262 | 265 | **1043** | **0** | 522 | 515 | 462 | 451 | **####** | **0** | 254 | 279 | 260 | 290 | **1083** | **0** |
| STK36 | *serine/threonine kinase 36* | 61 | 52 | 49 | 46 | **208** | **0** | 56 | 43 | 46 | 48 | **193** | **0** | 95 | 94 | 91 | 88 | **368** | **0** | 45 | 41 | 36 | 36 | **158** | **0** |
| LUZP1 | *leucine zipper protein 1* | 51 | 46 | 46 | 44 | **187** | **0** | 38 | 43 | 41 | 42 | 164 | 0.82 | 73 | 69 | 62 | 58 | **262** | **0** | 45 | 52 | 48 | 51 | 196 | 0.74 |
| STIP1 | *stress induced phosphoprotein 1* | 44 | 44 | 39 | 38 | **165** | **0** | 38 | 52 | 48 | 53 | 191 | 0.6 | 7 | 7 | 6 | 6 | 26 | 0.93 | 15 | 12 | 6 | 6 | 39 | 0.87 |
| HAUS8 | *HAUS augmin like complex subunit 8* | 38 | 34 | 33 | 33 | **138** | **0** | 47 | 36 | 41 | 43 | **167** | **0** | 47 | 41 | 41 | 36 | **165** | **0** | 53 | 61 | 64 | 53 | **231** | **0** |
| CCP110 | *centriolar coiled-coil protein 110* | 36 | 35 | 35 | 31 | **137** | **0** | 29 | 30 | 29 | 27 | **115** | **0** | 99 | 90 | 87 | 80 | **356** | **0** | 80 | 74 | 88 | 75 | **317** | **0** |
| CEP97 | *centrosomal protein 97* | 33 | 32 | 31 | 31 | **127** | **0** | 36 | 30 | 37 | 31 | **134** | **0** | 94 | 93 | 92 | 92 | **371** | **0** | 101 | 99 | 124 | 117 | **441** | **0** |
| MPHOSPH9 | *M-phase phosphoprotein 9* | 32 | 30 | 29 | 24 | **115** | **0** | 33 | 38 | 33 | 30 | **134** | **0** | 77 | 68 | 61 | 55 | **261** | **0** | 54 | 65 | 64 | 60 | **243** | **0** |
| PAN2 | *poly(A) specific ribonuclease subunit PAN2* | 30 | 28 | 27 | 26 | **111** | **0** | 21 | 29 | 25 | 38 | **113** | **0** | 86 | 80 | 74 | 68 | **308** | **0** | 82 | 74 | 64 | 73 | **293** | **0** |
| PDZD8 | *PDZ domain containing 8* | 30 | 28 | 27 | 26 | **111** | **0** | 20 | 32 | 31 | 23 | **106** | **0** | 10 | 7 | 7 | 6 | 30 | 0.59 | 17 | 19 | 16 | 11 | **63** | **0** |
| ROCK2 | *Rho associated coiled-coil containing protein kinase 2* | 27 | 26 | 24 | 24 | **101** | **0** | 18 | 17 | 21 | 19 | **75** | **0** | 68 | 59 | 55 | 52 | **234** | **0** | 52 | 60 | 73 | 68 | **253** | **0** |
| SIRT2 | *sirtuin 2* | 27 | 24 | 23 | 23 | **97** | **0** | 14 | 17 | 13 | 21 | **65** | **0** | 0 | 0 | 0 | 0 | 1 | na | 0 | 0 | 0 | 0 | 1 | na |
| HAUS2 | *HAUS augmin like complex subunit 2* | 24 | 24 | 23 | 23 | **94** | **0** | 29 | 25 | 26 | 31 | **111** | **0** | 31 | 30 | 25 | 24 | **110** | **0** | 28 | 32 | 31 | 38 | **129** | **0** |
| SMG7 | *SMG7 nonsense mediated mRNA decay factor* | 23 | 22 | 21 | 21 | **87** | **0** | 13 | 13 | 15 | 13 | 54 | 0.16 | 32 | 28 | 28 | 25 | **113** | **0** | 13 | 19 | 26 | 24 | **82** | **0** |
| UGGT1 | *UDP-glucose glycoprotein glucosyltransferase 1* | 21 | 21 | 20 | 15 | **77** | **0** | 9 | 13 | 13 | 9 | **44** | **0** | 7 | 6 | 5 | 4 | **22** | **0** | 3 | 2 | 4 | 4 | **13** | **0** |
| DLG5 | *discs large MAGUK scaffold protein 5* | 21 | 20 | 18 | 17 | **76** | **0** | 7 | 12 | 10 | 8 | **37** | **0** | 37 | 30 | 29 | 28 | **124** | **0** | 33 | 34 | 42 | 51 | **160** | **0** |
| TNRC6A | *trinucleotide repeat containing adaptor 6A* | 19 | 17 | 16 | 16 | **68** | **0** | 5 | 6 | 7 | 6 | 24 | 0.51 | 26 | 24 | 23 | 21 | **94** | **0** | 8 | 9 | 11 | 12 | 40 | 0.04 |
| PTPN14 | *protein tyrosine phosphatase non-receptor type 14* | 18 | 17 | 15 | 13 | **63** | **0** | 11 | 8 | 5 | 9 | **33** | **0** | 7 | 6 | 6 | 6 | **25** | **0** | 5 | 3 | 4 | 4 | **16** | **0** |
| CAMSAP3 | *calmodulin regulated spectrin associated protein family me* | 18 | 15 | 14 | 11 | **58** | **0** | 10 | 10 | 8 | 11 | **39** | **0.01** | 17 | 14 | 11 | 9 | **51** | **0.01** | 10 | 8 | 8 | 15 | 41 | 0.02 |
| KIAA1009 | *centrosomal protein 162* | 12 | 11 | 10 | 9 | **42** | **0** | 16 | 15 | 14 | 14 | **59** | **0** | 38 | 37 | 32 | 31 | **138** | **0** | 27 | 28 | 30 | 37 | **122** | **0** |
| DAPK3 | *death associated protein kinase 3* | 9 | 4 | 4 | 4 | **21** | **0** | 0 | 0 | 0 | 0 | 1 | na | 11 | 7 | 5 | 5 | **28** | **0** | 0 | 0 | 0 | 0 | 1 | na |
| SIPA1L3 | *signal induced proliferation associated 1 like 3* | 6 | 6 | 5 | 4 | **21** | **0** | 4 | 5 | 5 | 7 | **21** | **0** | 10 | 6 | 6 | 6 | **28** | **0** | 16 | 19 | 9 | 10 | **54** | **0** |
| ARHGAP29 | *Rho GTPase activating protein 29* | 6 | 6 | 5 | 4 | **21** | **0** | 3 | 5 | 4 | 7 | **19** | **0** | 7 | 6 | 5 | 5 | **23** | **0** | 8 | 11 | 9 | 12 | **40** | **0** |
| RC3H1 | *ring finger and CCCH-type domains 1* | 5 | 5 | 5 | 5 | **20** | **0** | 2 | 4 | 4 | 0 | **10** | **0.01** | 7 | 4 | 4 | 4 | **19** | **0** | 0 | 0 | 3 | 5 | 8 | 0.08 |
| PAN3 | *poly(A) specific ribonuclease subunit PAN3* | 7 | 4 | 4 | 4 | **19** | **0** | 5 | 0 | 0 | 5 | 10 | 0.07 | 15 | 14 | 14 | 11 | **54** | **0** | 23 | 30 | 21 | 31 | **105** | **0** |
| FYTTD1 | *forty-two-three domain containing 1* | 4 | 4 | 4 | 4 | **16** | **0** | 0 | 2 | 0 | 0 | 2 | 0.3 | 2 | 0 | 0 | 0 | 2 | 0.42 | 0 | 0 | 0 | 0 | 1 | na |
| PPP5C | *protein phosphatase 5 catalytic subunit* | 4 | 4 | 4 | 3 | **15** | **0** | 3 | 2 | 2 | 0 | 7 | 0.02 | 0 | 0 | 0 | 0 | 1 | na | 0 | 0 | 0 | 0 | 1 | na |
| SEPT15 | *Septin15* | 4 | 4 | 3 | 3 | **14** | **0** | 0 | 0 | 0 | 0 | 1 | na | 2 | 0 | 0 | 0 | 2 | 0.42 | 0 | 0 | 0 | 0 | 1 | na |
| CABLES1 | *Cdk5 and Abl enzyme substrate 1* | 10 | 9 | 8 | 8 | **35** | **0.01** | 5 | 4 | 3 | 5 | 17 | 0.06 | 5 | 4 | 2 | 2 | 13 | 0.29 | 3 | 2 | 4 | 0 | 9 | 0.2 |
| AIP | *aryl hydrocarbon receptor interacting protein* | 8 | 8 | 7 | 7 | **30** | **0.01** | 9 | 6 | 5 | 6 | 26 | 0.02 | 0 | 0 | 0 | 0 | 1 | na | 0 | 0 | 0 | 0 | 1 | na |
| POLR3A | *RNA polymerase III subunit A* | 6 | 3 | 3 | 3 | **15** | **0.01** | 2 | 0 | 4 | 3 | **9** | **0.01** | 2 | 2 | 2 | 0 | 6 | 0.13 | 0 | 0 | 0 | 0 | 1 | na |
| RICTOR | *RPTOR independent companion of MTOR complex 2* | 5 | 4 | 3 | 2 | **14** | **0.01** | 8 | 5 | 3 | 3 | **19** | **0** | 2 | 2 | 0 | 0 | 4 | 0.25 | 8 | 8 | 0 | 5 | **21** | **0** |
| RAC1 | *Rac family small GTPase 1* | 4 | 3 | 3 | 3 | **13** | **0.01** | 3 | 5 | 4 | 5 | **17** | **0** | 2 | 2 | 0 | 0 | 4 | 0.25 | 0 | 0 | 0 | 0 | 1 | na |
| SEC16A | *SEC16 homolog A, endoplasmic reticulum export factor* | 29 | 29 | 27 | 24 | 109 | 0.02 | 35 | 40 | 39 | 35 | **149** | **0** | 43 | 43 | 40 | 32 | **158** | **0** | 49 | 56 | 79 | 66 | **250** | **0** |
| WDR47 | *WD repeat domain 47* | 4 | 3 | 2 | 2 | 11 | 0.03 | 0 | 3 | 3 | 3 | **9** | **0** | 8 | 6 | 5 | 5 | **24** | **0** | 5 | 2 | 4 | 2 | **13** | **0.01** |
| KIF1B | *kinesin family member 1B* | 8 | 8 | 6 | 4 | 26 | 0.05 | 7 | 7 | 7 | 2 | **23** | **0.01** | 18 | 15 | 14 | 11 | **58** | **0** | 10 | 13 | 6 | 6 | **35** | **0.01** |
| APC | *APC regulator of WNT signaling pathway* | 9 | 8 | 8 | 7 | 32 | 0.07 | 9 | 17 | 16 | 11 | **53** | **0** | 17 | 14 | 12 | 11 | **54** | **0.01** | 15 | 25 | 29 | 31 | **100** | **0** |
| KIFAP3 | *kinesin associated protein 3* | 6 | 6 | 2 | 0 | 14 | 0.08 | 0 | 3 | 2 | 0 | 5 | 0.11 | 16 | 14 | 14 | 12 | **56** | **0** | 0 | 0 | 2 | 0 | 2 | 0.3 |
| CSPP1 | *centrosome and spindle pole associated protein 1* | 4 | 3 | 2 | 0 | 9 | 0.09 | 0 | 0 | 2 | 0 | 2 | 0.3 | 17 | 16 | 14 | 13 | **60** | **0** | 16 | 16 | 16 | 22 | **70** | **0** |
| USP54 | *ubiquitin specific peptidase 54* | 3 | 3 | 2 | 0 | 8 | 0.09 | 0 | 0 | 0 | 0 | 1 | na | 8 | 7 | 6 | 4 | **25** | **0** | 7 | 6 | 6 | 6 | **25** | **0** |
| MAST2 | *microtubule associated serine/threonine kinase 2* | 3 | 2 | 2 | 0 | 7 | 0.1 | 0 | 0 | 0 | 0 | 1 | na | 7 | 5 | 4 | 3 | **19** | **0** | 0 | 0 | 0 | 0 | 1 | na |
| MAP7D3 | *MAP7 domain containing 3* | 41 | 39 | 36 | 35 | 151 | 0.12 | 27 | 28 | 24 | 33 | 112 | 0.86 | 55 | 50 | 50 | 45 | **200** | **0** | 45 | 46 | 45 | 43 | 179 | 0.81 |
| SIPA1L1 | *signal induced proliferation associated 1 like 1* | 2 | 2 | 2 | 0 | 6 | 0.13 | 5 | 5 | 6 | 0 | **16** | **0** | 5 | 5 | 5 | 3 | **18** | **0** | 4 | 6 | 5 | 4 | **19** | **0** |
| KIF3B | *kinesin family member 3B* | 3 | 2 | 0 | 0 | 5 | 0.21 | 2 | 2 | 0 | 2 | 6 | 0.04 | 6 | 6 | 5 | 4 | **21** | **0** | 0 | 0 | 2 | 0 | 2 | 0.3 |
| OFD1 | *OFD1 centriole and centriolar satellite protein* | 23 | 23 | 20 | 19 | 85 | 0.25 | 24 | 26 | 22 | 21 | 93 | 0.05 | 36 | 32 | 31 | 31 | **130** | **0** | 43 | 39 | 56 | 53 | **191** | **0** |
| MFHAS1 | *malignant fibrous histiocytoma amplified sequence 1* | 2 | 2 | 0 | 0 | 4 | 0.25 | 2 | 2 | 3 | 2 | 9 | 0.02 | 10 | 10 | 8 | 6 | **34** | **0** | 5 | 5 | 4 | 0 | **14** | **0** |
| NHSL1 | *NHS like 1* | 5 | 4 | 3 | 2 | 14 | 0.28 | 4 | 4 | 5 | 5 | 18 | 0.06 | 9 | 9 | 9 | 6 | **33** | **0.01** | 10 | 16 | 13 | 11 | **50** | **0** |
| MTUS1 | *microtubule associated scaffold protein 1* | 15 | 11 | 11 | 11 | 48 | 0.41 | 8 | 12 | 12 | 7 | 39 | 0.46 | 31 | 29 | 26 | 26 | **112** | **0** | 24 | 24 | 28 | 23 | **99** | **0** |
| DDX20 | *DEAD-box helicase 20* | 13 | 9 | 9 | 8 | 39 | 0.41 | 21 | 26 | 22 | 26 | **95** | **0** | 19 | 18 | 17 | 17 | **71** | **0** | 42 | 47 | 53 | 61 | **203** | **0** |
| ROCK1 | *Rho associated coiled-coil containing protein kinase 1* | 6 | 6 | 6 | 6 | 24 | 0.41 | 4 | 3 | 3 | 2 | 12 | 0.46 | 42 | 34 | 31 | 31 | **138** | **0** | 34 | 30 | 43 | 35 | **142** | **0** |
| KIF3A | *kinesin family member 3A* | 3 | 3 | 3 | 2 | 11 | 0.41 | 5 | 0 | 0 | 2 | 7 | 0.28 | 11 | 10 | 8 | 6 | **35** | **0.01** | 4 | 3 | 0 | 3 | 10 | 0.18 |
| KIF27 | *kinesin family member 27* | 2 | 0 | 0 | 0 | 2 | 0.42 | 3 | 2 | 0 | 3 | 8 | 0.02 | 7 | 7 | 6 | 5 | **25** | **0** | 0 | 0 | 2 | 3 | 5 | 0.11 |
| KIF14 | *kinesin family member 14* | 16 | 14 | 14 | 13 | 57 | 0.6 | 22 | 16 | 26 | 17 | 81 | 0.05 | 24 | 20 | 20 | 19 | **83** | **0.01** | 42 | 38 | 45 | 50 | **175** | **0** |
| NAV1 | *neuron navigator 1* | 5 | 5 | 3 | 2 | 15 | 0.71 | 6 | 4 | 6 | 7 | 23 | 0.57 | 19 | 16 | 14 | 13 | **62** | **0** | 27 | 28 | 24 | 29 | **108** | **0** |
| KIAA1033 | *WASH complex subunit 4* | 16 | 15 | 14 | 13 | 58 | 0.72 | 13 | 8 | 17 | 12 | 50 | 0.6 | 32 | 28 | 28 | 27 | **115** | **0.01** | 41 | 44 | 36 | 49 | **170** | **0** |
| USP15 | *ubiquitin specific peptidase 15* | 50 | 50 | 49 | 45 | 194 | 0.93 | 43 | 30 | 28 | 29 | 130 | 0.9 | 276 | 248 | 243 | 241 | **####** | **0** | 260 | 266 | 235 | 225 | **986** | **0** |
| LMTK2 | *lemur tyrosine kinase 2* | 0 | 0 | 0 | 0 | 1 | na | 2 | 0 | 0 | 4 | 6 | 0.11 | 11 | 10 | 9 | 8 | **38** | **0** | 14 | 15 | 19 | 6 | **54** | **0** |
| KIAA0556 | *KIAA0556* | 0 | 0 | 0 | 0 | 1 | na | 0 | 0 | 0 | 0 | 1 | na | 6 | 5 | 5 | 4 | **20** | **0** | 3 | 6 | 5 | 9 | **23** | **0** |
| CCDC138 | *coiled-coil domain containing 138* | 0 | 0 | 0 | 0 | 1 | na | 0 | 0 | 0 | 0 | 1 | na | 3 | 3 | 3 | 3 | **12** | **0.01** | 5 | 6 | 6 | 8 | **25** | **0** |

**Supplemental Table S3: BioID high confidence proximity interactors for full-length ULK4 K39R mutant**

#### Supplemental Table S4

**FlagBirA*-ULK4**

| | | WT | | | | | | Kinase Domain | | | | | | | Armadillo Repeats | | | | | | |
|---|---|---|---|---|---|---|---|---|---|---|---|---|---|---|---|---|---|---|---|---|---|
| Gene Symbol | Gene Name | A1 | A2 | B1 | B2 | SUM | BFDR | A1 | A2 | B1 | B2 | SUM | BFDR | log2 KD/WT | A1 | A2 | B1 | B2 | SUM | BFDR | log2 ArmRep/WT |
| ULK4 | *unc-51 like kinase 4* | #### | #### | #### | #### | ##### | na | 751 | 815 | 854 | 908 | 3328 | na | -2.1 | 1136 | 1195 | 954 | 1119 | 4404 | na | -1.6 |
| birA | *E. coli biotin carboxylase* | #### | #### | #### | #### | ##### | na | 2843 | 2877 | 3427 | 3645 | ##### | na | -0.2 | 1629 | 1604 | 1250 | 1439 | 5922 | na | -1.3 |
| CAMSAP1 | *calmodulin regulated spectrin associated protein 1* | 293 | 291 | 289 | 284 | 1157 | 0 | 12 | 11 | 14 | 16 | 53 | 0.86 | -4.4 | 139 | 132 | 122 | 126 | 519 | 0 | -1.2 |
| USP15 | *ubiquitin specific peptidase 15* | 50 | 50 | 49 | 45 | 194 | 0.93 | 58 | 56 | 66 | 50 | 230 | 0.89 | 0.2 | 22 | 22 | 19 | 17 | 80 | 0.89 | -1.3 |
| CEP97 | *centrosomal protein 97* | 33 | 32 | 31 | 31 | 127 | 0 | 8 | 9 | 6 | 7 | 30 | 0.59 | -2.1 | 11 | 12 | 6 | 8 | 37 | 0.46 | -1.8 |
| STK36 | *serine/threonine kinase 36* | 61 | 52 | 49 | 46 | 208 | 0 | 17 | 17 | 15 | 17 | 66 | 0 | -1.7 | 0 | 0 | 0 | 0 | 1 | na | -7.7 |
| CCP110 | *centriolar coiled-coil protein 110* | 36 | 35 | 35 | 31 | 137 | 0 | 3 | 0 | 3 | 6 | 12 | 0.28 | -3.5 | 6 | 6 | 4 | 6 | 22 | 0.08 | -2.6 |
| PAN2 | *poly(A) specific ribonuclease subunit PAN2* | 30 | 28 | 27 | 26 | 111 | 0 | 0 | 0 | 0 | 0 | 1 | na | -6.8 | 25 | 25 | 23 | 24 | 97 | 0 | -0.2 |
| LUZP1 | *leucine zipper protein 1* | 51 | 46 | 46 | 44 | 187 | 0 | 35 | 37 | 33 | 40 | 145 | 0.54 | -0.4 | 3 | 5 | 8 | 8 | 24 | 0.89 | -3.0 |
| MPHOSPH9 | *M-phase phosphoprotein 9* | 32 | 30 | 29 | 24 | 115 | 0 | 0 | 2 | 4 | 6 | 12 | 0.22 | -3.3 | 20 | 19 | 20 | 21 | 80 | 0 | -0.5 |
| ROCK2 | *Rho associated coiled-coil containing protein kinase 2* | 27 | 26 | 24 | 24 | 101 | 0 | 2 | 0 | 0 | 2 | 4 | 0.66 | -4.7 | 0 | 0 | 0 | 0 | 1 | na | -6.7 |
| MAP7D3 | *MAP7 domain containing 3* | 41 | 39 | 36 | 35 | 151 | 0.12 | 25 | 25 | 29 | 26 | 105 | 0.69 | -0.5 | 3 | 0 | 4 | 4 | 11 | 0.89 | -3.8 |
| HAUS8 | *HAUS augmin like complex subunit 8* | 38 | 34 | 33 | 33 | 138 | 0 | 35 | 38 | 46 | 41 | 160 | 0 | 0.2 | 43 | 50 | 39 | 38 | 170 | 0 | 0.3 |
| SEC16A | *SEC16 homolog A, endoplasmic reticulum export facto* | 29 | 29 | 27 | 24 | 109 | 0.02 | 45 | 42 | 42 | 38 | 167 | 0 | 0.6 | 33 | 31 | 26 | 28 | 118 | 0 | 0.1 |
| KIAA1009 | *centrosomal protein 162* | 12 | 11 | 10 | 9 | 42 | 0 | 0 | 0 | 0 | 0 | 1 | na | -5.4 | 4 | 5 | 3 | 4 | 16 | 0 | -1.4 |
| ROCK1 | *Rho associated coiled-coil containing protein kinase 1* | 6 | 6 | 6 | 6 | 24 | 0.41 | 0 | 0 | 0 | 0 | 1 | na | -4.6 | 0 | 0 | 0 | 0 | 1 | na | -4.6 |
| OFD1 | *OFD1 centriole and centriolar satellite protein* | 23 | 23 | 20 | 19 | 85 | 0.25 | 15 | 19 | 18 | 14 | 66 | 0.58 | -0.4 | 71 | 66 | 66 | 60 | 263 | 0 | 1.6 |
| DLG5 | *discs large MAGUK scaffold protein 5* | 21 | 20 | 18 | 17 | 76 | 0 | 14 | 9 | 11 | 13 | 47 | 0 | -0.7 | 11 | 9 | 6 | 5 | 31 | 0.01 | -1.3 |
| SMG7 | *SMG7 nonsense mediated mRNA decay factor* | 23 | 22 | 21 | 21 | 87 | 0 | 15 | 17 | 15 | 16 | 63 | 0.01 | -0.5 | 0 | 6 | 6 | 6 | 18 | 0.69 | -2.3 |
| MTUS1 | *microtubule associated scaffold protein 1* | 15 | 11 | 11 | 11 | 48 | 0.41 | 3 | 0 | 2 | 4 | 9 | 0.74 | -2.4 | 0 | 0 | 0 | 0 | 1 | na | -5.6 |
| HAUS2 | *HAUS augmin like complex subunit 2* | 24 | 24 | 23 | 23 | 94 | 0 | 23 | 23 | 23 | 23 | 92 | 0 | 0.0 | 33 | 43 | 28 | 49 | 153 | 0 | 0.7 |
| TNRC6A | *trinucleotide repeat containing adaptor 6A* | 19 | 17 | 16 | 16 | 68 | 0 | 6 | 4 | 7 | 8 | 25 | 0.5 | -1.4 | 13 | 7 | 4 | 7 | 31 | 0.27 | -1.1 |
| DDX20 | *DEAD-box helicase 20* | 13 | 9 | 9 | 8 | 39 | 0.41 | 5 | 5 | 6 | 6 | 22 | 0.64 | -0.8 | 39 | 37 | 31 | 33 | 140 | 0 | 1.8 |
| NAV1 | *neuron navigator 1* | 5 | 5 | 3 | 2 | 15 | 0.71 | 0 | 0 | 0 | 2 | 2 | 0.71 | -2.9 | 11 | 11 | 9 | 7 | 38 | 0.12 | 1.3 |
| CSPP1 | *centrosome and spindle pole associated protein 1* | 4 | 3 | 2 | 0 | 9 | 0.09 | 0 | 0 | 0 | 0 | 1 | na | -3.2 | 5 | 5 | 0 | 5 | 1 | 0.04 | -3.2 |
| KIF1B | *kinesin family member 1B* | 8 | 8 | 6 | 4 | 26 | 0.05 | 5 | 0 | 0 | 5 | 10 | 0.24 | -1.4 | 0 | 0 | 0 | 0 | 1 | na | -4.7 |
| KIFAP3 | *kinesin associated protein 3* | 6 | 6 | 2 | 0 | 14 | 0.08 | 0 | 0 | 0 | 0 | 1 | na | -3.8 | 0 | 0 | 0 | 0 | 1 | na | -3.8 |
| PAN3 | *poly(A) specific ribonuclease subunit PAN3* | 7 | 4 | 4 | 4 | 19 | 0 | 0 | 0 | 0 | 2 | 2 | 0.39 | -3.2 | 0 | 2 | 0 | 0 | 1 | 0.39 | -4.2 |
| LMTK2 | *lemur tyrosine kinase 2* | 0 | 0 | 0 | 0 | 1 | na | 0 | 0 | 0 | 0 | 1 | na | 0.0 | 0 | 0 | 0 | 0 | 1 | na | 0.0 |
| MFHAS1 | *malignant fibrous histiocytoma amplified sequence 1* | 2 | 2 | 0 | 0 | 4 | 0.25 | 0 | 0 | 0 | 0 | 1 | na | -2.0 | 0 | 0 | 0 | 0 | 1 | na | -2.0 |
| DAPK3 | *death associated protein kinase 3* | 9 | 4 | 4 | 4 | 21 | 0 | 0 | 0 | 0 | 0 | 1 | na | -4.4 | 0 | 0 | 0 | 0 | 1 | na | -4.4 |
| SIPA1L3 | *signal induced proliferation associated 1 like 3* | 6 | 6 | 5 | 4 | 21 | 0 | 0 | 4 | 2 | 5 | 11 | 0.08 | -0.9 | 0 | 4 | 0 | 0 | 4 | 0.32 | -2.4 |
| PTPN14 | *protein tyrosine phosphatase non-receptor type 14* | 18 | 17 | 15 | 13 | 63 | 0 | 6 | 3 | 9 | 10 | 28 | 0 | -1.2 | 0 | 0 | 0 | 0 | 1 | na | -6.0 |
| USP54 | *ubiquitin specific peptidase 54* | 3 | 3 | 2 | 0 | 8 | 0.09 | 0 | 0 | 0 | 0 | 1 | na | -3.0 | 0 | 2 | 0 | 0 | 2 | 0.39 | -2.0 |
| KIF27 | *kinesin family member 27* | 2 | 0 | 0 | 0 | 2 | 0.42 | 0 | 0 | 0 | 2 | 2 | 0.39 | 0.0 | 0 | 0 | 0 | 0 | 1 | na | -1.0 |
| WDR47 | *WD repeat domain 47* | 4 | 3 | 2 | 2 | 11 | 0.03 | 0 | 0 | 0 | 0 | 1 | na | -3.5 | 0 | 0 | 0 | 0 | 1 | na | -3.5 |
| ARHGAP29 | *Rho GTPase activating protein 29* | 6 | 6 | 5 | 4 | 21 | 0 | 3 | 2 | 0 | 0 | 5 | 0.19 | -2.1 | 0 | 2 | 0 | 0 | 2 | 0.39 | -3.4 |
| UGGT1 | *UDP-glucose glycoprotein glucosyltransferase 1* | 21 | 21 | 20 | 15 | 77 | 0 | 5 | 5 | 5 | 3 | 18 | 0 | -2.1 | 4 | 3 | 2 | 0 | 9 | 0.08 | -3.1 |
| KIF3B | *kinesin family member 3B* | 3 | 2 | 0 | 0 | 5 | 0.21 | 0 | 0 | 0 | 0 | 1 | na | -2.3 | 0 | 0 | 0 | 0 | 1 | na | -2.3 |
| KIAA0556 | *KIAA0556* | 0 | 0 | 0 | 0 | 1 | na | 0 | 0 | 0 | 0 | 1 | na | 0.0 | 0 | 0 | 0 | 0 | 1 | na | 0.0 |
| RC3H1 | *ring finger and CCCH-type domains 1* | 5 | 5 | 5 | 5 | 20 | 0 | 0 | 0 | 0 | 0 | 1 | na | -4.3 | 0 | 0 | 0 | 0 | 1 | na | -4.3 |
| MAST2 | *microtubule associated serine/threonine kinase 2* | 3 | 2 | 2 | 0 | 7 | 0.1 | 0 | 0 | 0 | 0 | 1 | na | -2.8 | 0 | 0 | 0 | 0 | 1 | na | -2.8 |
| SIPA1L1 | *signal induced proliferation associated 1 like 1* | 2 | 2 | 2 | 0 | 6 | 0.13 | 0 | 0 | 0 | 0 | 1 | na | -2.6 | 0 | 0 | 0 | 0 | 1 | na | -2.6 |
| KIAA1033 | *WASH complex subunit 4* | 16 | 15 | 14 | 13 | 58 | 0.72 | 7 | 11 | 9 | 7 | 34 | 0.73 | -0.8 | 16 | 15 | 9 | 17 | 57 | 0.61 | 0.0 |
| KIF14 | *kinesin family member 14* | 16 | 14 | 14 | 13 | 57 | 0.6 | 6 | 5 | 7 | 6 | 24 | 0.75 | -1.2 | 9 | 9 | 6 | 8 | 32 | 0.71 | -0.8 |
| APC | *APC regulator of WNT signaling pathway* | 9 | 8 | 8 | 7 | 32 | 0.07 | 7 | 9 | 0 | 3 | 19 | 0.13 | -0.8 | 15 | 16 | 7 | 12 | 50 | 0.01 | 0.6 |
| CAMSAP3 | *calmodulin regulated spectrin associated protein fam* | 18 | 15 | 14 | 11 | 58 | 0 | 2 | 2 | 0 | 0 | 4 | 0.5 | -3.9 | 0 | 0 | 0 | 0 | 1 | na | -5.9 |
| KIF3A | *kinesin family member 3A* | 3 | 3 | 3 | 2 | 11 | 0.41 | 0 | 0 | 0 | 0 | 1 | na | -3.5 | 2 | 2 | 0 | 0 | 1 | 0.48 | -3.5 |
| NHSL1 | *NHS like 1* | 5 | 4 | 3 | 2 | 14 | 0.28 | 0 | 0 | 0 | 0 | 1 | na | -3.8 | 0 | 0 | 0 | 0 | 1 | na | -3.8 |
| CCDC138 | *coiled-coil domain containing 138* | 0 | 0 | 0 | 0 | 1 | na | 0 | 0 | 0 | 0 | 1 | na | 0.0 | 20 | 21 | 15 | 20 | 76 | 0 | 6.2 |
| POLR3A | *RNA polymerase III subunit A* | 6 | 3 | 3 | 3 | 15 | 0.01 | 0 | 3 | 4 | 0 | 7 | 0.16 | -1.1 | 7 | 6 | 5 | 0 | 18 | 0.04 | 0.3 |
| RICTOR | *RPTOR independent companion of MTOR complex 2* | 5 | 4 | 3 | 2 | 14 | 0.01 | 0 | 0 | 0 | 0 | 1 | na | -3.8 | 0 | 0 | 0 | 0 | 1 | na | -3.8 |
| RAC1 | *Rac family small GTPase 1* | 4 | 3 | 3 | 3 | 13 | 0.01 | 0 | 0 | 0 | 0 | 1 | na | -3.7 | 3 | 4 | 5 | 4 | 16 | 0 | 0.3 |
| CABLES1 | *Cdk5 and Abl enzyme substrate 1* | 10 | 9 | 8 | 8 | 35 | 0.01 | 6 | 5 | 8 | 6 | 25 | 0.02 | -0.5 | 0 | 0 | 0 | 0 | 1 | na | -5.1 |
| FYTTD1 | *forty-two-three domain containing 1* | 4 | 4 | 4 | 4 | 16 | 0 | 3 | 3 | 0 | 4 | 10 | 0.07 | -0.7 | 0 | 3 | 2 | 2 | 7 | 0.1 | -1.2 |
| SEPT15 | *Septin15* | 4 | 4 | 3 | 3 | 14 | 0 | 0 | 0 | 0 | 0 | 1 | na | -3.8 | 0 | 0 | 0 | 0 | 1 | na | -3.8 |
| PDZD8 | *PDZ domain containing 8* | 30 | 28 | 27 | 26 | 111 | 0 | 7 | 6 | 7 | 5 | 25 | 0.6 | -2.2 | 0 | 0 | 0 | 0 | 1 | na | -6.8 |
| STIP1 | *stress induced phosphoprotein 1* | 44 | 44 | 39 | 38 | 165 | 0 | 42 | 37 | 38 | 42 | 159 | 0.01 | -0.1 | 58 | 62 | 55 | 52 | 227 | 0 | 0.5 |
| SIRT2 | *sirtuin 2* | 27 | 24 | 23 | 23 | 97 | 0 | 8 | 13 | 13 | 9 | 43 | 0 | -1.2 | 0 | 0 | 0 | 0 | 1 | na | -6.6 |
| PPP5C | *protein phosphatase 5 catalytic subunit* | 4 | 4 | 4 | 3 | 15 | 0 | 0 | 2 | 0 | 0 | 2 | 0.39 | -2.9 | 0 | 2 | 0 | 0 | 1 | 0.39 | -3.9 |
| AIP | *aryl hydrocarbon receptor interacting protein* | 8 | 8 | 7 | 7 | 30 | 0.01 | 4 | 5 | 3 | 4 | 16 | 0.11 | -0.9 | 6 | 8 | 11 | 8 | 33 | 0 | 0.1 |

**ULK4-BirA*Flag**

| | WT | | | | | | Kinase Domain | | | | | | | Armadillo Repeats | | | | | | |
|---|---|---|---|---|---|---|---|---|---|---|---|---|---|---|---|---|---|---|---|---|
| Gene Symbol | A1 | A2 | B1 | B2 | SUM | BFDR | A1 | A2 | B1 | B2 | SUM | BFDR | log2 KD/WT | A1 | A2 | B1 | B2 | SUM | BFDR | log2 ArmRep/WT |
| ULK4 | #### | #### | #### | #### | #### | na | 696 | 670 | 680 | 877 | 2923 | na | -1.6 | 145 | 146 | 154 | 137 | 582 | na | -4.0 |
| birA | #### | #### | #### | #### | #### | na | 2450 | 2389 | 2389 | 2506 | 9734 | na | 0.1 | 232 | 219 | 263 | 222 | 936 | na | -3.3 |
| CAMSAP1 | 522 | 515 | 462 | 451 | #### | 0 | 16 | 20 | 16 | 14 | 66 | 0.85 | -4.9 | 175 | 176 | 193 | 162 | 706 | 0 | -1.5 |
| USP15 | 276 | 248 | 243 | 241 | #### | 0 | 53 | 60 | 45 | 48 | 206 | 0.89 | -2.3 | 45 | 48 | 66 | 58 | 217 | 0.89 | -2.2 |
| CEP97 | 94 | 93 | 92 | 92 | 371 | 0 | 117 | 122 | 111 | 116 | 466 | 0 | 0.3 | 0 | 3 | 3 | 6 | 12 | 0.68 | -5.0 |
| STK36 | 95 | 94 | 91 | 88 | 368 | 0 | 28 | 25 | 36 | 31 | 120 | 0 | -8.5 | 0 | 0 | 0 | 0 | 1 | na | -8.5 |
| CCP110 | 99 | 90 | 87 | 80 | 356 | 0 | 83 | 81 | 89 | 90 | 343 | 0 | -0.1 | 0 | 0 | 0 | 0 | 1 | na | -8.5 |
| PAN2 | 86 | 80 | 74 | 68 | 308 | 0 | 0 | 0 | 0 | 0 | 1 | na | -8.3 | 18 | 21 | 17 | 20 | 76 | 0.01 | -2.0 |
| LUZP1 | 73 | 69 | 62 | 58 | 262 | 0 | 31 | 28 | 25 | 25 | 109 | 0.75 | -1.3 | 8 | 6 | 6 | 4 | 24 | 0.89 | -3.4 |
| MPHOSPH9 | 77 | 68 | 61 | 55 | 261 | 0 | 12 | 11 | 14 | 15 | 52 | 0 | -2.3 | 0 | 2 | 4 | 6 | 12 | 0.22 | -4.4 |
| ROCK2 | 68 | 59 | 55 | 52 | 234 | 0 | 0 | 0 | 0 | 0 | 1 | na | -7.9 | 0 | 2 | 0 | 2 | 4 | 0.66 | -5.9 |
| MAP7D3 | 55 | 50 | 50 | 45 | 200 | 0 | 25 | 23 | 20 | 20 | 88 | 0.76 | -1.2 | 7 | 6 | 7 | 8 | 28 | 0.86 | -2.8 |
| HAUS8 | 47 | 41 | 41 | 36 | 165 | 0 | 55 | 47 | 46 | 38 | 186 | 0 | 0.2 | 25 | 26 | 23 | 28 | 102 | 0.53 | -0.7 |
| SEC16A | 43 | 43 | 40 | 32 | 158 | 0 | 169 | 153 | 170 | 169 | 661 | 0 | 2.1 | 8 | 7 | 12 | 9 | 36 | 0.77 | -2.1 |
| KIAA1009 | 38 | 37 | 32 | 31 | 138 | 0 | 0 | 0 | 0 | 0 | 1 | na | -7.1 | 4 | 0 | 4 | 4 | 12 | 0.05 | -3.5 |
| ROCK1 | 42 | 34 | 31 | 31 | 138 | 0 | 0 | 0 | 0 | 0 | 1 | na | -7.1 | 0 | 0 | 0 | 0 | 1 | na | -7.1 |
| OFD1 | 36 | 32 | 31 | 31 | 130 | 0 | 26 | 23 | 22 | 25 | 96 | 0.02 | -0.4 | 34 | 31 | 39 | 29 | 133 | 0 | 0.0 |
| DLG5 | 37 | 30 | 29 | 28 | 124 | 0 | 20 | 21 | 23 | 17 | 81 | 0 | -0.6 | 6 | 6 | 4 | 8 | 24 | 0.02 | -2.4 |
| SMG7 | 32 | 28 | 28 | 25 | 113 | 0 | 20 | 23 | 24 | 18 | 85 | 0 | -0.4 | 0 | 2 | 5 | 4 | 11 | 0.72 | -3.4 |
| MTUS1 | 31 | 29 | 26 | 26 | 112 | 0 | 2 | 3 | 0 | 4 | 9 | 0.74 | -3.6 | 0 | 0 | 0 | 0 | 1 | na | -6.8 |
| HAUS2 | 31 | 30 | 25 | 24 | 110 | 0 | 34 | 37 | 32 | 34 | 137 | 0 | 0.3 | 26 | 22 | 22 | 24 | 94 | 0 | -0.2 |
| TNRC6A | 26 | 24 | 23 | 21 | 94 | 0 | 14 | 11 | 14 | 6 | 45 | 0.06 | -1.1 | 6 | 12 | 5 | 10 | 33 | 0.21 | -1.5 |
| DDX20 | 19 | 18 | 17 | 17 | 71 | 0 | 6 | 13 | 9 | 13 | 41 | 0.21 | -0.8 | 28 | 28 | 20 | 18 | 94 | 0 | 0.4 |
| NAV1 | 19 | 16 | 14 | 13 | 62 | 0 | 0 | 0 | 2 | 2 | 4 | 0.7 | -4.0 | 5 | 4 | 5 | 8 | 22 | 0.55 | -1.5 |
| CSPP1 | 17 | 16 | 14 | 13 | 60 | 0 | 0 | 0 | 0 | 0 | 1 | na | -5.9 | 0 | 0 | 0 | 0 | 1 | na | -5.9 |
| KIF1B | 18 | 15 | 14 | 11 | 58 | 0 | 7 | 6 | 3 | 2 | 18 | 0.11 | -1.7 | 0 | 0 | 0 | 0 | 1 | na | -5.9 |
| KIFAP3 | 16 | 14 | 14 | 12 | 56 | 0 | 0 | 0 | 0 | 0 | 1 | na | -5.8 | 0 | 0 | 0 | 2 | 2 | 0.39 | -4.8 |
| PAN3 | 15 | 14 | 14 | 11 | 54 | 0 | 0 | 0 | 0 | 0 | 1 | na | -5.8 | 0 | 0 | 0 | 0 | 1 | na | -5.8 |
| LMTK2 | 11 | 10 | 9 | 8 | 38 | 0 | 0 | 0 | 0 | 0 | 1 | na | -5.2 | 0 | 0 | 0 | 0 | 1 | na | -5.2 |
| MFHAS1 | 10 | 10 | 8 | 6 | 34 | 0 | 0 | 0 | 0 | 0 | 1 | na | -5.1 | 0 | 0 | 2 | 0 | 2 | 0.39 | -4.1 |
| DAPK3 | 11 | 7 | 5 | 5 | 28 | 0 | 0 | 0 | 0 | 0 | 1 | na | -4.8 | 0 | 0 | 0 | 0 | 1 | na | -4.8 |
| SIPA1L3 | 10 | 6 | 6 | 6 | 28 | 0 | 10 | 5 | 8 | 11 | 34 | 0 | 0.3 | 0 | 0 | 0 | 0 | 1 | na | -4.8 |
| PTPN14 | 7 | 6 | 6 | 6 | 25 | 0 | 7 | 15 | 13 | 16 | 51 | 0 | 1.0 | 0 | 0 | 0 | 0 | 1 | na | -4.6 |
| USP54 | 8 | 7 | 6 | 4 | 25 | 0 | 0 | 2 | 0 | 0 | 2 | 0.39 | -3.6 | 0 | 0 | 0 | 0 | 1 | na | -4.6 |
| KIF27 | 7 | 7 | 6 | 5 | 25 | 0 | 0 | 0 | 0 | 0 | 1 | na | -4.6 | 0 | 0 | 0 | 0 | 1 | na | -4.6 |
| WDR47 | 8 | 6 | 5 | 5 | 24 | 0 | 0 | 0 | 0 | 0 | 1 | na | -4.6 | 0 | 0 | 0 | 0 | 1 | na | -4.6 |
| ARHGAP29 | 7 | 6 | 5 | 5 | 23 | 0 | 2 | 0 | 2 | 4 | 8 | 0.09 | -1.5 | 0 | 0 | 0 | 0 | 1 | na | -4.5 |
| UGGT1 | 7 | 6 | 5 | 4 | 22 | 0 | 3 | 3 | 0 | 4 | 10 | 0.07 | -1.1 | 0 | 0 | 0 | 0 | 1 | na | -4.5 |
| KIF3B | 6 | 6 | 5 | 4 | 21 | 0 | 0 | 0 | 0 | 0 | 1 | na | -4.4 | 0 | 0 | 0 | 0 | 1 | na | -4.4 |
| KIAA0556 | 6 | 5 | 5 | 4 | 20 | 0 | 0 | 0 | 0 | 0 | 1 | na | -4.3 | 0 | 0 | 0 | 0 | 1 | na | -4.3 |
| RC3H1 | 7 | 4 | 4 | 4 | 19 | 0 | 15 | 12 | 11 | 10 | 48 | 0 | 1.3 | 0 | 0 | 0 | 0 | 1 | na | -4.2 |
| MAST2 | 7 | 5 | 4 | 3 | 19 | 0 | 7 | 8 | 8 | 7 | 30 | 0 | 0.7 | 0 | 0 | 0 | 0 | 1 | na | -4.2 |
| SIPA1L1 | 5 | 5 | 5 | 3 | 18 | 0 | 0 | 3 | 0 | 0 | 3 | 0.33 | -2.6 | 0 | 0 | 0 | 0 | 1 | na | -4.2 |
| KIAA1033 | 32 | 28 | 28 | 27 | 115 | 0.01 | 7 | 8 | 6 | 10 | 31 | 0.74 | -1.9 | 6 | 8 | 8 | 8 | 30 | 0.76 | -1.9 |
| KIF14 | 24 | 20 | 20 | 19 | 83 | 0.01 | 6 | 6 | 6 | 5 | 23 | 0.75 | -1.9 | 6 | 6 | 7 | 7 | 26 | 0.74 | -1.7 |
| APC | 17 | 14 | 12 | 11 | 54 | 0.01 | 5 | 6 | 4 | 7 | 22 | 0.11 | -1.3 | 3 | 4 | 6 | 5 | 18 | 0.13 | -1.6 |
| CAMSAP3 | 17 | 14 | 11 | 9 | 51 | 0.01 | 7 | 13 | 12 | 10 | 42 | 0.01 | -0.3 | 0 | 0 | 0 | 0 | 1 | na | -5.7 |
| KIF3A | 11 | 10 | 8 | 6 | 35 | 0.01 | 0 | 0 | 0 | 0 | 1 | na | -5.1 | 3 | 0 | 5 | 6 | 14 | 0.12 | -1.3 |
| NHSL1 | 9 | 9 | 9 | 6 | 33 | 0.01 | 0 | 3 | 0 | 3 | 6 | 0.38 | -2.5 | 0 | 0 | 0 | 0 | 1 | na | -5.0 |
| CCDC138 | 3 | 3 | 3 | 3 | 12 | 0.01 | 0 | 0 | 0 | 0 | 1 | na | -3.6 | 5 | 6 | 10 | 11 | 32 | 0 | 1.4 |
| POLR3A | 2 | 2 | 2 | 0 | 6 | 0.13 | 11 | 13 | 14 | 9 | 47 | 0 | 3.0 | 0 | 0 | 0 | 0 | 1 | na | -2.6 |
| RICTOR | 2 | 2 | 0 | 0 | 4 | 0.25 | 7 | 3 | 9 | 4 | 23 | 0 | 2.5 | 0 | 0 | 0 | 0 | 1 | na | -2.0 |
| RAC1 | 2 | 2 | 0 | 0 | 4 | 0.25 | 2 | 4 | 0 | 2 | 8 | 0.09 | 1.0 | 4 | 3 | 5 | 4 | 16 | 0 | 2.0 |
| CABLES1 | 5 | 4 | 2 | 2 | 13 | 0.29 | 2 | 0 | 6 | 3 | 11 | 0.24 | -0.2 | 0 | 0 | 2 | 0 | 2 | 0.52 | -2.7 |
| FYTTD1 | 2 | 0 | 0 | 0 | 2 | 0.42 | 4 | 3 | 3 | 2 | 12 | 0.01 | 2.6 | 2 | 0 | 0 | 0 | 2 | 0.39 | 0.0 |
| SEPT15 | 2 | 0 | 0 | 0 | 2 | 0.42 | 0 | 0 | 0 | 0 | 1 | na | -1.0 | 0 | 0 | 0 | 0 | 1 | na | -1.0 |
| PDZD8 | 10 | 7 | 7 | 6 | 30 | 0.59 | 29 | 32 | 30 | 33 | 124 | 0 | 2.0 | 0 | 0 | 0 | 0 | 1 | na | -4.9 |
| STIP1 | 7 | 7 | 6 | 6 | 26 | 0.93 | 16 | 17 | 12 | 12 | 57 | 0.82 | 1.1 | 19 | 14 | 18 | 18 | 69 | 0.81 | 1.4 |
| SIRT2 | 0 | 0 | 0 | 0 | 1 | na | 14 | 15 | 16 | 12 | 57 | 0 | 5.8 | 0 | 0 | 0 | 0 | 1 | na | 0.0 |
| PPP5C | 0 | 0 | 0 | 0 | 1 | na | 0 | 0 | 0 | 0 | 1 | na | 0.0 | 0 | 0 | 0 | 0 | 1 | na | 0.0 |
| AIP | 0 | 0 | 0 | 0 | 1 | na | 4 | 0 | 0 | 0 | 4 | 0.47 | 2.0 | 0 | 0 | 0 | 0 | 1 | na | 0.0 |

Supplemental Table S4: BioID high confidence proximity interactors for the Armadillo and Kinase domain of ULK4
